# Drought and recovery in barley: key gene networks and retrotransposon response

**DOI:** 10.1101/2023.03.05.531133

**Authors:** Maitry Paul, Jaakko Tanskanen, Marko Jääskeläinen, Wei Chang, Ahan Dalal, Menachem Moshelion, Alan H. Schulman

**Affiliations:** HiLIFE Institute of Biotechnology, University of Helsinki, Helsinki, Finland; Viikki Plant Science Centre (ViPS), University of Helsinki, Helsinki, Finland; Natural Resources Institute Finland (LUKE), Helsinki, Finland; The Robert H Smith Institute of Plant Sciences and Genetics in Agriculture, The Hebrew University of Jerusalem, Rehovot, Israel

**Keywords:** Barley (*Hordeum vulgare* L.), drought, resilience, recovery, gene expression analysis, *BARE* retrotransposon, autophagy

## Abstract

- During drought, plants close their stomata at a critical soil water content (SWC), together with diverse physiological, developmental, and biochemical responses.
- Using precision-phenotyping lysimeters, we imposed pre-flowering drought on four barley varieties (Arvo, Golden Promise, Hankkija 673 and Morex) and followed their physiological responses. For Golden Promise, we carried out RNA-seq on leaf transcripts before and during drought, and during recovery, also examining retrotransposon *BARE1* expression. Transcriptional data were subjected to network analysis.
- The varieties differed by their critical SWC, Hankkija 673 responding at the highest and Golden Promise at the lowest. Pathways connected to drought and salinity response were strongly upregulated during drought; pathways connected to growth and development were strongly downregulated. During recovery, growth and development pathways were upregulated; altogether 117 networked genes involved in ubiquitin-mediated autophagy were downregulated. The differential response to SWC suggests adaptation to distinct rainfall patterns.
- We identified several strongly differentially expressed genes not earlier associated with drought response in barley. *BARE1* transcription is strongly transcriptionally upregulated by drought and downregulated during recovery unequally between the investigated cultivars. The downregulation of networked autophagy genes suggests a role for autophagy in drought response; its importance to resilience should be further investigated.

## INTRODUCTION

Optimization of the balance between carbon fixed and water lost by transpiration is a central problem for plants. In drought, the challenge to water homeostasis and increased water potential generates a response by the plant at a critical soil water content (SWC), leading to stomatal closure, even in daylight, to reduce water loss. The associated dehydration and its consequences are commonly referred to as drought stress. Drought response involves mechanisms and signaling cascades leading to stomatal closure (Merilo *et al.*, 2015) and to physiological and metabolic responses (Jogawat *et al.*, 2021). Mechanisms to counter dehydration and cellular damage include accumulation of osmolytes to mediate osmotic adjustment (Hildebrandt, 2018) and enzymatic and non-enzymatic scavenging of excess reactive oxygen species (ROS) (Das & Roychoudhury, 2014), as well as changes in the chloroplast proteome (Chen *et al.*, 2021).

Two contrasting responses to water limitation have been described: isohydric and anisohydric (Sade *et al.*, 2012). Isohydric implies maintenance of constant water potential; many physiological parameters are involved (Scharwies & Dinneny, 2019). Barley has been shown to be diurnally anisohydric (Tardieu & Simonneau, 1998). The terms also have been applied broadly for drought response strategy (Sade *et al.*, 2012), isohydric plants conservating water, closing stomata at a relatively high SWC, sacrificing carbon fixation but delaying plant dehydration. This strategy may reduce risk from early droughts in climates were the probability of precipitation increases during the growing season. An anisohydric plant delays stomatal closure and continues fixing carbon, a strategy consistent with environments having terminal droughts, or with those where dry periods are short and show little seasonal variation.

Studies in Arabidopsis indicate that induction of autophagy is needed for drought tolerance (Avin-Wittenberg, 2019). Autophagy is an evolutionarily conserved mechanism for recycling damaged proteins and cellular organelles by transport to the vacuoles or lysosomes for degradation (Chen *et al.*, 2021). Rewatering at the end of drought may result in the resumption of normal diurnal stomatal opening, recovery of photosynthesis, and the resumption of growth. The occurrence, degree, and rate of recovery strongly depends both on the intensity and duration of drought and on the species; recovery has relatively little studied on its own (Xu *et al.*, 2010; Lawas *et al.*, 2019; Chen *et al.*, 2021; Qi *et al.*, 2021). Recovery involves not only the ABA signaling pathway (Cao *et al.*, 2021), but also strigolactone (Visentin *et al.*, 2020).

Drought is also well known to induce transcription of transposable elements, particularly retrotransposons (RLX; Wicker *et al.* (2007). The RLXs have been shown to be transcriptionally induced by stresses ranging from ionizing and UV radiation to drought and heavy metals (Kimura *et al.*, 2001; Grandbastien *et al.*, 2005; Ramallo *et al.*, 2008; Makarevitch *et al.*, 2015; Galindo-Gonzalez *et al.*, 2017). For barley (*Hordum vulgare* L.) RLX, earlier studies indicated that *BARE1* transcripts and translational products accumulated under drought (Jääskeläinen *et al.*, 2013) and appeared linked to the abscisic acid (ABA) signaling pathway through the ABA-response elements (ABREs) present in the *BARE1* promoter (Jääskeläinen *et al.*, 2013).

Optimization of drought response for an annual crop such as barley is related to the climatic zone for which the plant is bred. In some zones, the probability of drought may be greater early in the growing season (e.g., Nordic conditions, before flowering); in others, the probability of drought increases during seed maturation (Mediterranean basin, as terminal drought) (Rollins *et al.*, 2013; Tao *et al.*, 2017). Here, we focus on pre-flowering drought in four barley cultivars shown in pilot experiments to respond differentially to SWC (Dalal et al., in prep). For one of them (Golden Promise), the goal was to carry out transcriptional analyses for leaves for both genes and the *BARE1* RLX at physiologically defined time points before, during, and after drought. Gene expression data was then input to network analysis to understand the activated pathways and possible connections to RLX stress response.

## MATERIALS AND METHODS

### Plant material

Barley cultivars Arvo, Golden Promise (“GP”), Hankkija 673 (“H673”) and Morex was selected from the diversity set of the ERA-NET SusCrop Climbar Project. GP is the standard genotype for transformation in barley (Schreiber *et al.*, 2019), whereas preliminary experiments indicated that Arvo and H673 have high and low stomatal conductance respectively (Dalal et al., in prep.). Morex is the established reference genome assembly in barley (Beier *et al.*, 2017; Mascher *et al.*, 2017).

### Growth conditions

Plants were grown during the winter season in the iCORE functional-phenotyping greenhouse (https://plantscience.agri.huji.ac.il/icore-center) under natural light, with moderate temperature control, which closely reflects the natural external environment (Halperin *et al.*, 2017; Galkin *et al.*, 2018) (Fig. 1a). There, drought experiments were carried out with a randomized-block design on a gravimetric functional phenotyping platform (Dalal *et al.*, 2020; Gosa SC, 2022) (Plantarray, PA 3.0, PlantDitech Ltd., Yavne, Israel), which allows simultaneous standardized, control of drought treatment with continuous monitoring of multiple physiological parameters. The platform comprises temperature-compensated load cells (lysimeters) for simultaneous and continuous gravimetric monitoring of water relations in the soil–plant–atmosphere continuum (SPAC) under dynamic environmental conditions (Halperin *et al.*, 2017).

**Fig. 1.**
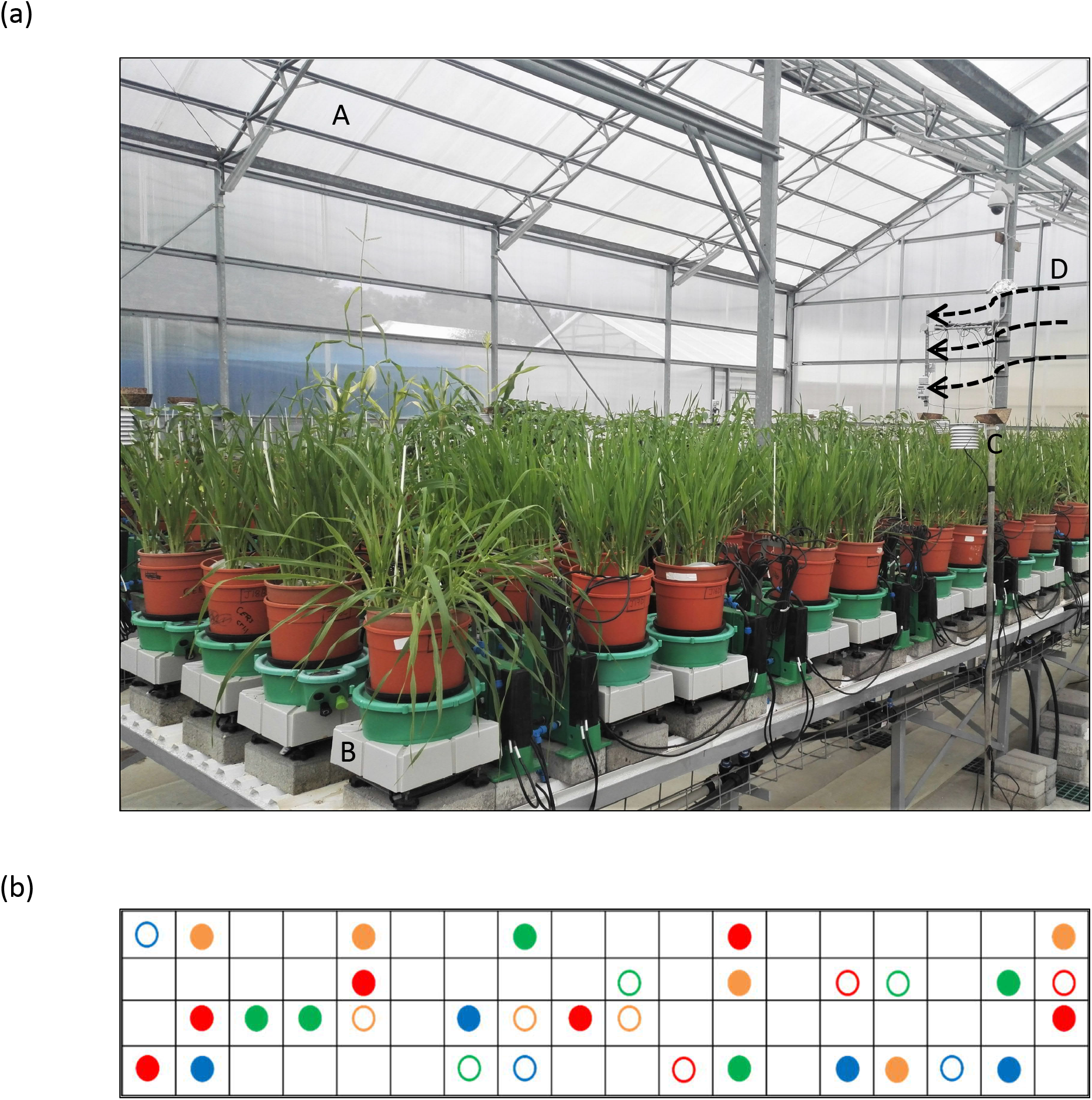
Experimental setup. (a) Lysimeters of gravimetric functional phenotyping platform, comprising 72 lysimeter platforms with soil probes and four weather stations. The setup features: (A) minimally controlled greenhouse; (B) lysimeter platform; (C) weather station; (D) cooling vent. (b) Table diagram for sample pots distributed according to a randomized block design. Filled circles are drought treatments; empty circles are well-watered (controls). Arvo, orange; GP, red; H673, green; Morex, blue.

Seeds were sown in germination trays, then transplanted at the 3-4-leaf stage (two weeks old) to pots. Before transplanting, seedling roots were carefully washed to remove the original soil, planted into potting soil (“Bental 11”, Tuff Marom Golan, Israel; Dalal *et al.* (2020) four per 3.9-litre pot, then placed onto lysimeters (Dalal *et al.*, 2019; Gosa SC, 2022). Each cultivar was grown in four to six biological replicates for the drought treatment and in three biological replicates for the controls, which were maintained under well-irrigated conditions (Fig. 1b). The plants were grown at 18–25°C, 20–35% relative humidity, and 10h light /14h dark. Temperature, humidity, and photosynthetically active radiation (PAR) were monitored as described earlier (Dalal *et al.*, 2020; Gosa SC, 2022). The vapor pressure deficit (VPD) ranged between 0.2 to 4.0 kPa in the greenhouse and represented typical weather fluctuations during winters in Central Israel (Fig. 2b).

**Fig. 2.**
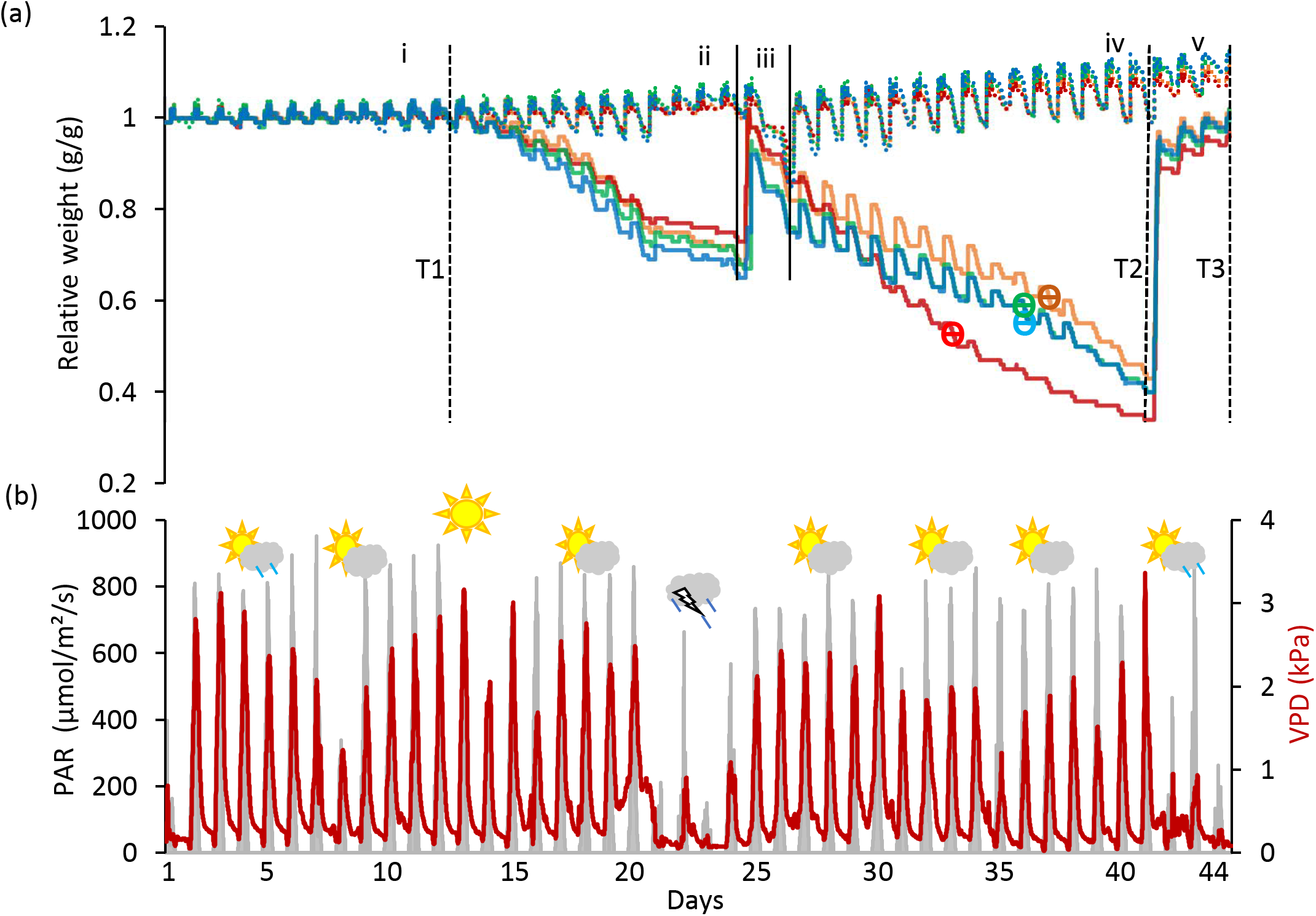
Key system parameters during drought and recovery. (a) Relative change in system weight of all control and drought-treated pots. The experiment is divided into five phases: (i) well-watered; (ii) Dry Phase I; (iii) Rewatering I; (iv) Dry Phase II; (v) Rewatering II. Drought samples, solid lines; well-irrigated controls, dashed. Cultivars are: Arvo, orange; Golden Promise (GP), red; Hankkija 673 (H673), green; Morex, blue. Day of ϴ crit indicated by ϴ on the cultivars’ curves. Time points (T) for collection of RNA from leaf samples are: T1, last day of well-watered; T2, last day of drought; T3, last day of recovery. (b) Photosynthetic Active Radiation (PAR) and Vapor Pressure Deficit (VPD) over the course of the experiment. The weather symbols depict conditions outdoors during this time.

### Experimental design

To mimic the onset of a drought, the experiment was divided into five phases on the lysimeter platform: (i) well-watered; (ii) Dry Phase I (iii); Rewatering I (iv); Dry Phase II; (v) Rewatering II (Fig. 2a). For the well-watered phase (first 12 days) and for the controls throughout, plants daily received three nighttime irrigations at 3-hour intervals to reach field capacity. The volumetric water content (VWC) was calculated with the SPAC analytical software embedded in the Plantarray system (Fig. 3). In Dry Phase I, the system was set to irrigate droughted plants to 80% of their own previous day’s transpiration, so that these would be subjected to gradual water stress from day 13 to day 24, followed by rewatering (Rewatering I) on the night of day 25. Dry Phase II extended from day 26 to day 41 with the same irrigation parameters as Dry Phase I, until the time when the droughted plants reached <100g daily transpiration (around 20 to 30% of controls) (Fig. 4d). Rewatering II began on day 42. Daily night irrigation was given to all plants until day 44. At that point, several irrigation cycles were used to moisten the soil and to ensure a laterally uniform soil water content (SWC) in the pots. VPD data were continuously monitored (Fig. 2b).

**Fig. 3.**
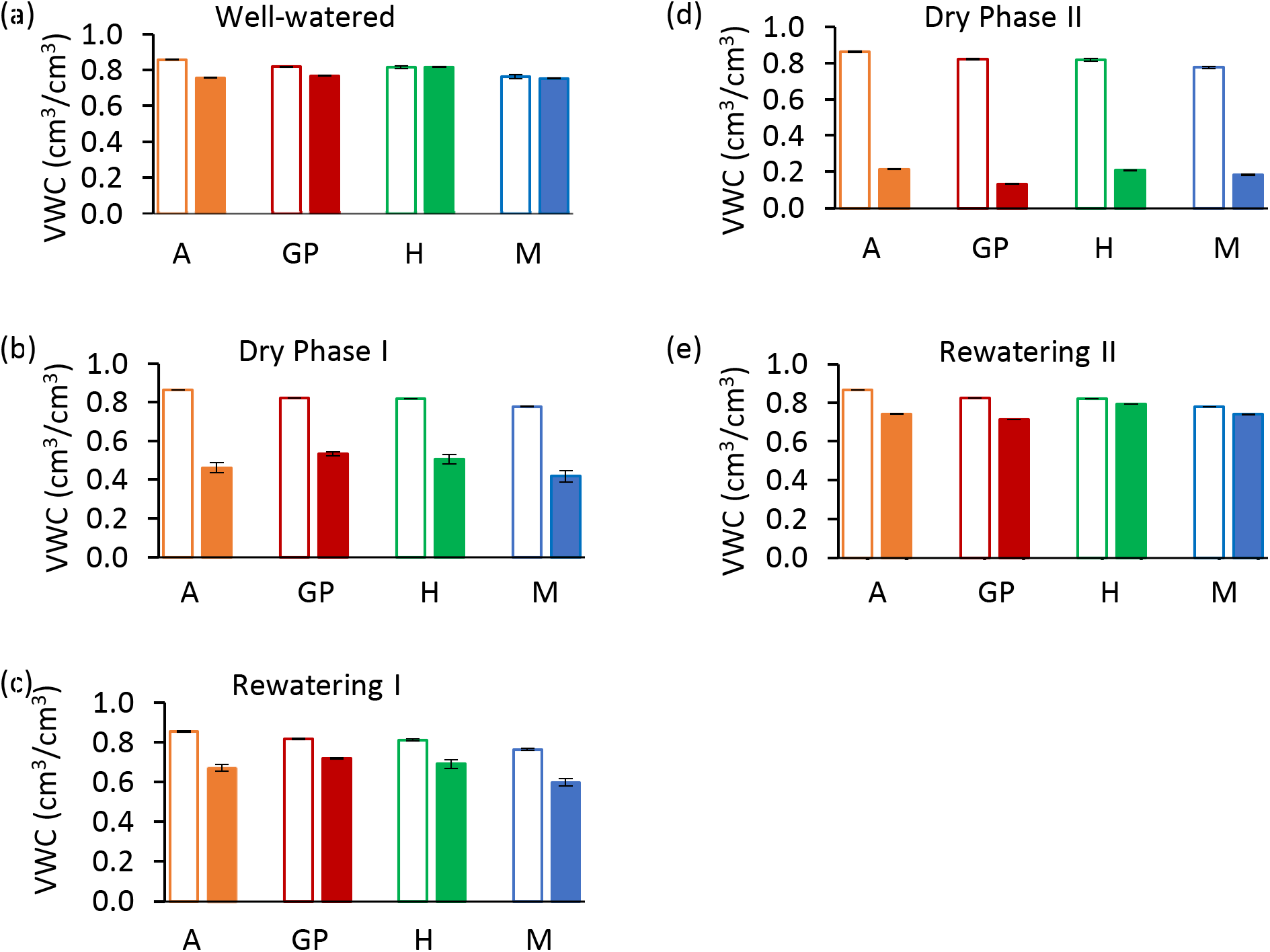
Calculated soil volumetric water content (VWC) during five phases of the experiment. (a) Day 12, last day of well-watered (Phase i). (b) Day 24, last day of Dry Phase I. (c) Day 25, last day of Rewatering I. (d) Day 41, Dry Phase II. (e) Day 44, last day of Rewatering II. Solid bars represent drought samples; empty bars represent controls. Arvo (A), orange; red, Golden Promise (GP), green; Hankkija 673 (H); blue, Morex (M).

**Fig 4.**
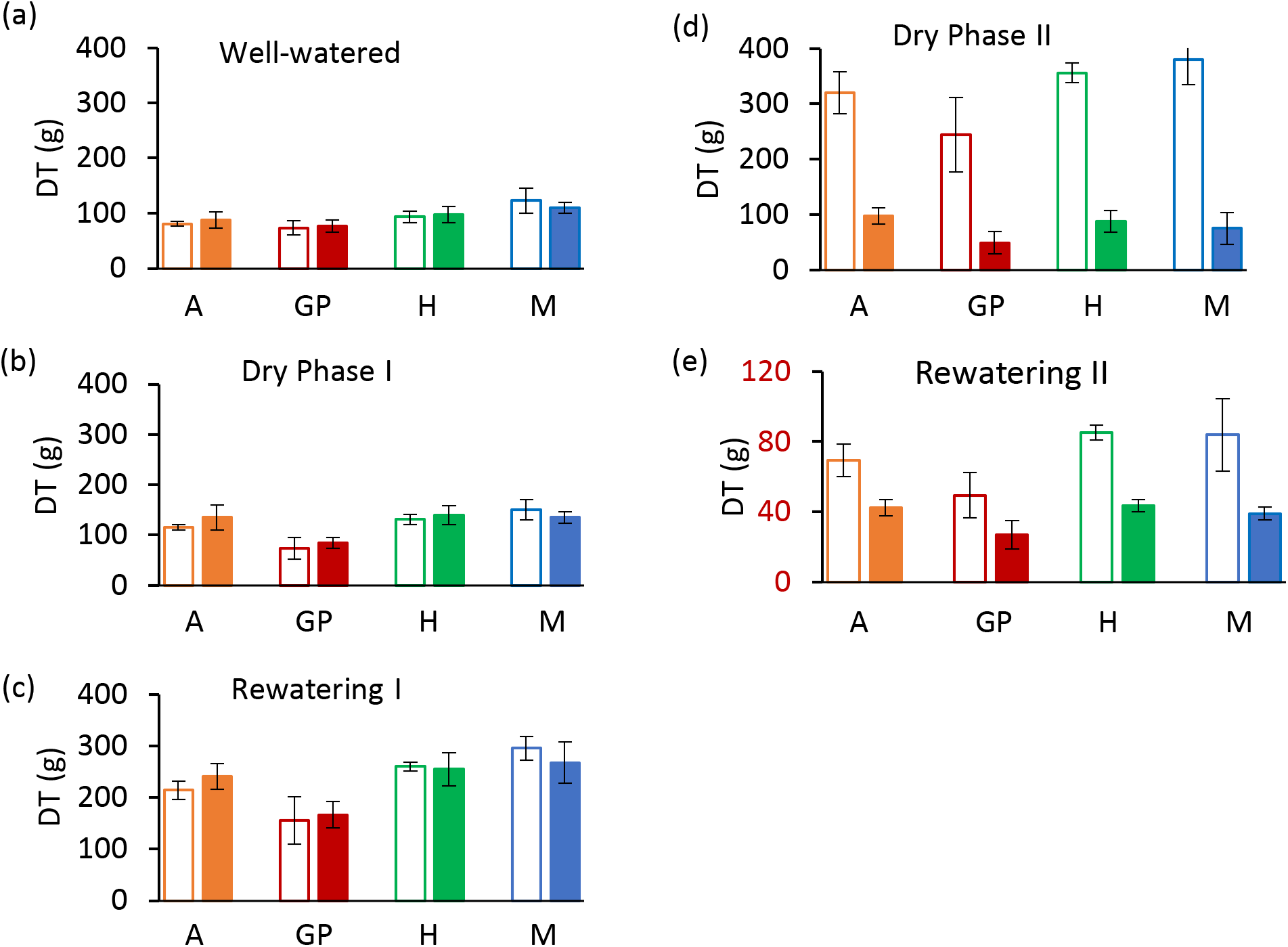
Daily Transpiration (DT) at the end of each experimental phase. (a) Day 12, last day of well-watered (Phase i). (b) Day 24, last day of Dry Phase I. (c) Day 25, last day of Rewatering I. (d) Day 41, Dry Phase II. (e) Day 44, last day of Rewatering II. Solid bars represent drought samples, empty bars controls. Orange represents Arvo (A); red, Golden Promise (GP); green, Hankkija 673 (H); blue, Morex (M).

### Sampling and isolation of RNA

Second leaves from the youngest tillers were collected for RNA analysis: Timepoint 1 (T1) on the last day of the well-watered phase (day 12); T2, on the last day of Dry Phase II (day 41); T3, last day of Rewatering II (day 44). Well-watered control samples at the same growth stages were collected at T2 and T3. For RNA extraction, leaf samples were collected in liquid N_2_ and later ground and stored in TRIzol reagent (Invitrogen). Total RNA was extracted from the leaf samples with the Direct-zol RNA Miniprep Plus kit according to the manufacturer’s (Zymo Research Europe GmbH) guidelines. RNA (1 µg) from each sample was treated with Ambion® DNase I (RNase-free; Applied Biosystems) in a 20 µl reaction mixture for 90 min at 37 °C to remove remaining DNA. An RNeasy MinElute Cleanup Kit (Qiagen) was used to purify and concentrate the RNA from the DNase treatments. RNA quantity and quality were checked both spectrophotometrically (NANODROP-2000 spectrophotometer, Thermo Scientific) and by agarose gel electrophoresis.

### qPCR analysis

Reverse transcription to cDNA was carried out in reactions containing 200 ng of total RNA, 1 µl Invitrogen™ SuperScript® IV Reverse Transcriptase (RT) (Thermo Fisher Scientific), 1 µl 10 mM dNTPs (Thermo Scientific) and 0.5 µl 10 µM primer E1820 (AAGCAGTGGTAACAACGCAGAGTACT_30_NA) as the oligo(dT) primer (Chang *et al.*, 2013). For qPCR, 10 µl reactions comprised 0.5 µl cDNA as template, 5 µl SsoAdvanced™ Universal SYBR® Green Supermix, 0.1 µl each forward and reverse primer, and 4.3 µl water. Reactions were run on a CFX 384 Real-Time PCR detection system (Bio-Rad) in three technical replicates as follows: activation for 2 min at 94°C; 40 cycles comprising denaturation for 30 sec at 94°C, annealing for 30 sec at 56°C, and extension for 1 min at 72°C. Relative quantification of the cDNA product was made by the 2**^- ΔΔCT^** method (Livak & Schmittgen, 2001). All primers used for qPCR are listed in Supporting Information Table S1. The amplification efficiency of all the primers has been checked; R^2^ was from 0.97-0.99.

### RNA-seq and network analysis

Three sets of biological replicates from each timepoint and their corresponding controls were taken from *cv.* GP for RNA-seq analysis. Libraries were constructed using the TruSeq Stranded Total RNA kit with plant cytoplasmic and chloroplast ribosomal RNA removal (Illumina); adaptors were TruSeq adaptors. FastQC (Andrews, 2010) was used to check the quality control metrics of the raw reads and to trim the RNA-seq data. Adapter sequences were removed with Trimmomatic (Bolger *et al.*, 2014). The STAR alignment method (Dobin *et al.*, 2013) served to map sequence reads to the GP genome (GPV1.48, Ensembl Plants). Next, to quantify transcript expression, the Salmon tool was applied (Patro *et al.*, 2017). Differentially expressed genes (DEGs) and PCA plots were obtained with DESeq2 (Love *et al.*, 2014; Hong *et al.*, 2020). MDS plots were used to verify the approximate expression differences between sample replicates.

From the clean reads, those with pADJ value <0.05 for their fold-change were run in g: Profiler for functional enrichment analysis and to obtain a GMT file for Cytoscape (Reimand *et al.*, 2019). Cytoscape_v3.8.2 was then used to develop network clusters for biological pathways. The parameters used were: FDR q value cutoff =1, node cutoff 0.5, edge cutoff 0.375, and significance threshold for functional enrichment *p* ≤ 0.01.

## RESULTS

### Whole-plant water relations over the course of drought and recovery

Four barley varieties (Arvo, GP, H673, and Morex), which in preliminary experiments differed in their water use strategies, were placed on lysimeters and measured through the four experimental phases. Transpiration and plant weight were followed continuously. The soil water content (SWC) at which the transpiration rate drops in response to drought (ϴ critical point; ϴcrit) was calculated from the data.

#### Daily transpiration

Analysis of daily transpiration (DT) of the four varieties on the Plantarray platform revealed variation in their transpiration both between the varieties (Figs. 4, Supporting Information Fig. S1) and day-by-day (Fig. 5a, S1a-c). All plants were exposed to similar changes in PAR and VPD simultaneously (Fig. 1), which were monitored throughout the experiment (Fig. 2b); from Day 21 to 23, the cloudy and rainy weather outside the minimally controlled greenhouse reduced the PAR and VPD within, lowering DT. For the well-watered control plants of all varieties, DT gradually increased along with plant size over the course of the experiment, closely matched by the droughted plants, which then diverged following the onset of Dry Phase II. The overall DT of Morex is the highest, followed by H673, Arvo, and GP. The DT of droughted Morex plants diverged from controls by Day 20 (Fig. S1c) and H673 on Day 25 (Fig. S1b), whereas Arvo and GP only from Day 30 (Fig. 5a, S1a). A complete return to control whole-plant transpiration was not observed for any of the varieties before termination of the experiment on Day 44; all plants showed low DT between Days 42 and 44, linked to low VPD and PAR.

**Fig. 5.**
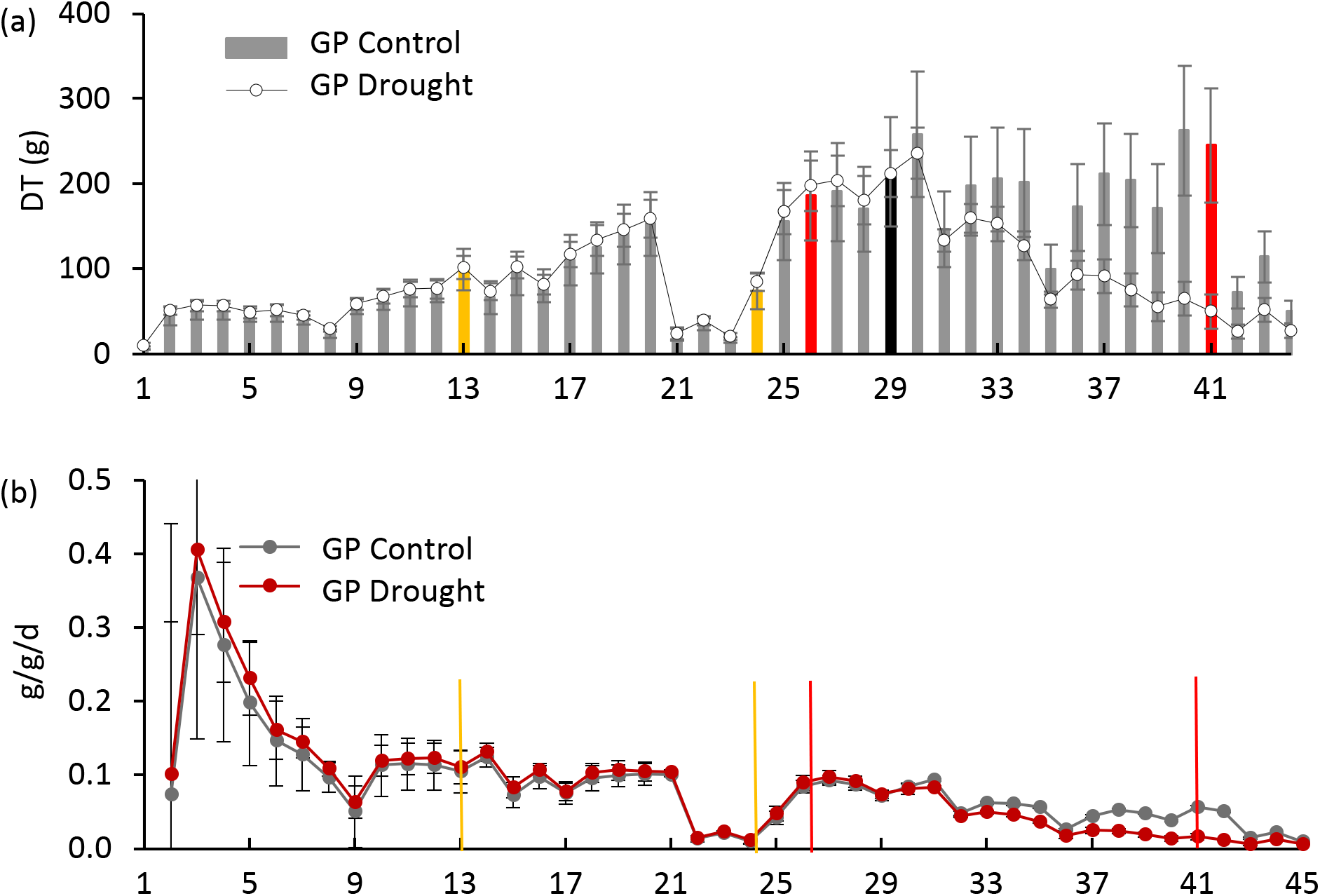
Daily transpiration and plant weight over the experimental period for Golden Promise (GP). (a) Daily transpiration (DT). The black bar represents the divergence of drought from control plants. (b) Daily specific change in plant weight (g/g/d). Yellow bars represent the start and end of Dry Phase I, red bars show the start and end of Dry Phase II.

### Effect of drought on the plant growth and transpiration

Changes in net plant weight as a measure of growth was calculated from the Plantarray system as before (Halperin *et al.*, 2017). The high rate of growth observed for all varieties between days 2 and 9 (Fig. 5b, S2 a-c) corresponds to the peak in PAR during the same period (Fig. 2b). The control plants showed similar trends in growth rates throughout the experiment. The droughted plants of all four varieties displayed specific growth rates lower than their well-watered controls during drought, first in Morex (Day 23), then Arvo (Day 29), then on Day 30 both H673 and GP (Fig. S2, 5b). The relative effect of drought on the growth rate of the four varieties is most apparent when the rate of change is considered.

The ϴ_crit_ was calculated for all varieties (Fig. 6) based on midday whole plant transpiration (Halperin *et al.*, 2017). The transpiration rates of all four varieties remained constant up to a VWC of 0.55 and then declined, with the VWC at the ϴ_crit_ differing by variety. Arvo, H673, and Morex display similar initial transpiration rates, whereas GP was about 30% lower. The H673 ϴ_crit_ occurred at 0.53 VWC on Day 36, Arvo at 0.52 on Day 37, Morex at 0.46 on Day 36, and GP at 0.42 already on Day 33. Following stomatal closure at ϴ_crit_, weight (water) loss preceded at different rates among the varieties. Although GP had the lowest initial transpiration rate (Fig. 6), it declined the fastest, whereas Arvo, with a high transpiration rate, declined the slowest.

**Fig. 6.**
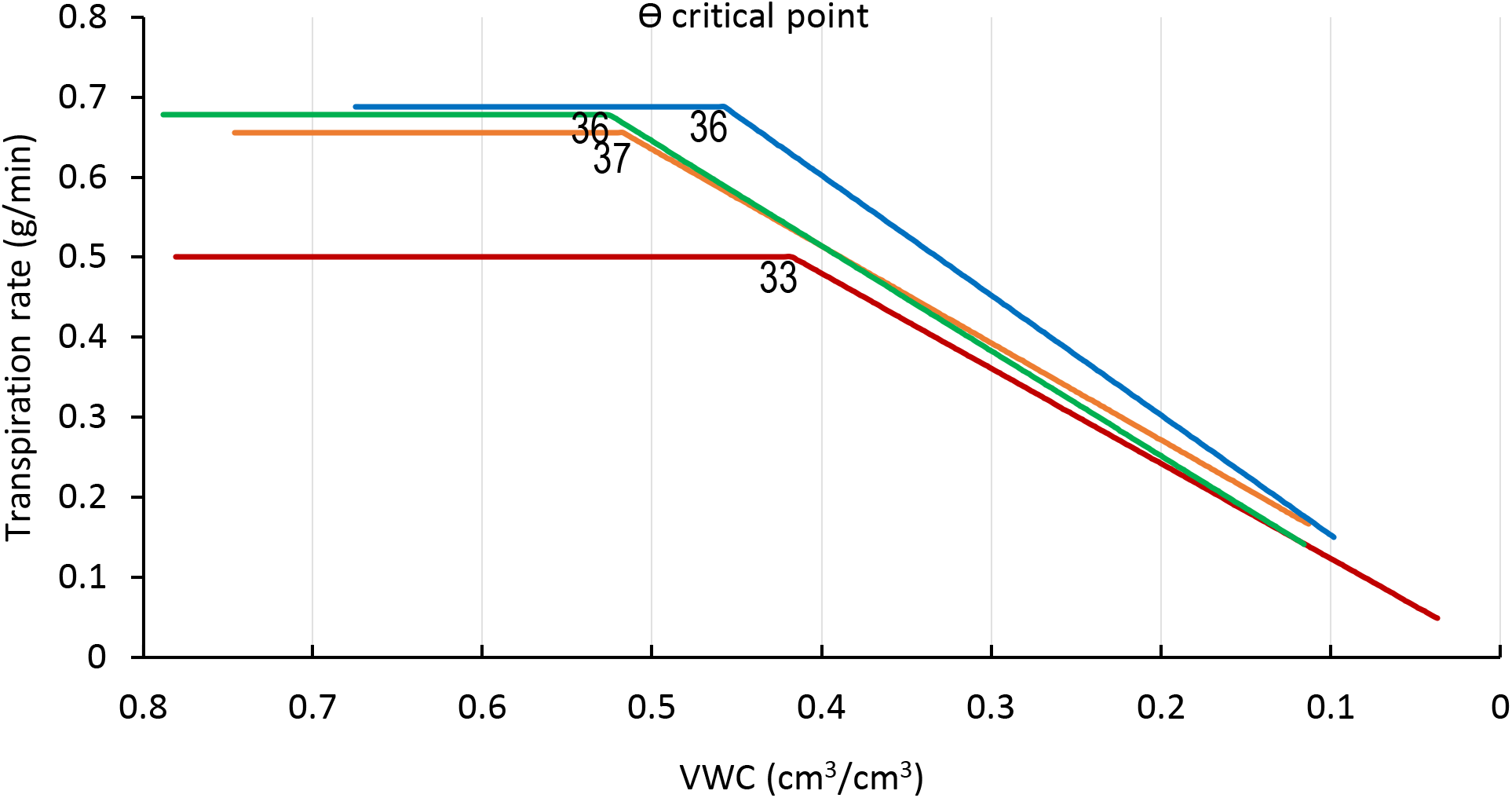
ϴ critical point (ϴ_crit_), for midday whole plant transpiration versus calculated VWC. The inflection points in the fitted transpiration lines indicate the ϴ_crit_. Arvo, orange; Golden Promise, red; Hankkija 673 (H673), green; Morex, blue. Day on which the ϴ_crit_ occurred is indicated by a number adjacent to the inflection point.

### Gene expression analysis

An average of 33 million reads per sample were generated by RNA-seq, which showed the expression of approximately 27,000 genes. To check the quality of the gene expression, DESeq2 dispersion estimates were made to reflect the variance in gene expression for a given mean value. Dispersion estimates were made and values adjusted as the basis for significance testing (Fig. S3). Principal components analysis (PCA) of differential gene expression showed an alignment of all groups of samples along PC1 and a narrow dispersion of all control groups along PC2 (Fig. S4). Movement of the expression pattern during the course of the experiment shows a shift within PC2 during drought and then a return to near the position of the control groups for the recovery samples (Fig. S4), consistent with ongo08ing physiological recovery. Of the genes detected, those differentially expressed (DEGs) with adjusted *p* < 0.01 include 611 that were over-expressed and 643 under-expressed in drought compared to the well-watered controls samples taken on the same day, while 697 were over-expressed and 282 under-expressed during recovery compared with the controls on the same day (Table S2-S5).

RNA-seq reads were annotated by reference to the cv. Morex genome (release 2), transcript expression quantified, and DEGs identified. In the drought samples, the most highly over-expressed gene compared to the well-watered control was for glutathione s-transferase (Table 1), with 3-million-fold more expression than control; in the samples from plants undergoing recovery, this gene dropped by 8.6-fold compared with drought. Altogether, 44 genes showed more than 100-fold overexpression and 218 genes over 10-fold overexpression compared to controls (Table S2). The most underexpressed gene was for a cytochrome P450 family protein (Table 2), with around 3000-fold less expression than control. Conversely, among the DEGs in recovery compared to drought, this gene showed the third-highest overexpression, 195-fold (Table S7). Altogether, 9 genes showed more than 100-fold underexpression and 53 genes over 10-fold underexpression (Table S3).

**Table 1.** Top 20 overexpressed genes, drought phase.

| Ensembl_gene_id | Description | log2 FC<sup>1</sup> | padj<sup>2</sup> |
| --- | --- | --- | --- |
| HORVU.MOREX.r2.5HG0441700 | glutathione s-transferase | 21.49 | 5.6E-07 |
| HORVU.MOREX.r2.6HG0456270 | Linalool synthase | 13.20 | 9.4E-05 |
| HORVU.MOREX.r2.6HG0516720 | Dehydrin | 11.76 | 9.4E-18 |
| HORVU.MOREX.r2.5HG0395660 | U-box domain containing protein | 10.67 | 6.2E-03 |
| HORVU.MOREX.r2.5HG0429440 | Dehydrin | 10.31 | 3.4E-05 |
| HORVU.MOREX.r2HG0053260 | NOT FOUND | 10.17 | 2.1E-03 |
| HORVU.MOREX.r2.3HG0242240 | Coiled-coil domain-containing protein 18 | 9.89 | 7.6E-12 |
| HORVU.MOREX.r2.3HG0184850 | Trypsin inhibitor | 9.44 | 5.5E-09 |
| HORVU.MOREX.r2.4HG0328360 | Branched-chain-amino-acid aminotransferase | 9.38 | 5.6E-04 |
| HORVU.MOREX.r2.3HG0186590 | Basic 7S globulin 2 | 9.28 | 2.1E-04 |
| HORVU.MOREX.r2.7HG0593880 | cysteine-rich/transmembrane domain A-like protein | 9.19 | 2.3E-03 |
| HORVU.MOREX.r2.3HG0201230 | DWNN domain | 9.13 | 1.9E-04 |
| HORVU.MOREX.r2.7HG0620080 | Carbonic anhydrase | 9.00 | 2.7E-03 |
| HORVU.MOREX.r2.2HG0149830 | Caleosin | 8.39 | 4.3E-05 |
| HORVU.MOREX.r2.3HG0184880 | Trypsin inhibitor | 8.38 | 6.9E-11 |
| HORVU.MOREX.r2.3HG0245240 | Disulfide isomerase | 8.35 | 4.6E-07 |
| HORVU.MOREX.r2.5HG0356770 | Germin-like protein | 8.09 | 8.1E-04 |
| HORVU.MOREX.r2.5HG0402770 | basic helix-loop-helix (bHLH) DNA-binding superfamily protein | 8.04 | 2.7E-05 |
| HORVU.MOREX.r2.2HG0149820 | Caleosin | 7.75 | 9.0E-03 |
| HORVU.MOREX.r2.6HG0512380 | 2-oxoglutarate (2OG) and Fe (II)-dependent oxygenase superfamily protein | 7.61 | 1.7E-03 |

<sup>1</sup>FC= fold-change
<sup>2</sup>padj= adjusted *p* value

**Table 2.** Top 20 underexpressed genes, drought phase.

| Ensembl_gene_id | Description | log2 FC<sup>1</sup> | padj<sup>2</sup> |
| --- | --- | --- | --- |
| HORVU.MOREX.r2.3HG0191820 | Cytochrome P450 family protein | -11.71 | 6.0E-08 |
| HORVU.MOREX.r2.3HG0268430 | Basic helix-loop-helix (BHLH)<br>family transcription factor | -9.82 | 3.1E-03 |
| HORVU.MOREX.r2.4HG0346330 | Cortical cell-delineating protein | -9.46 | 2.6E-09 |
| HORVU.MOREX.r2.4HG0346340 | Cortical cell-delineating protein | -9.28 | 4.7E-08 |
| HORVU.MOREX.r2.2HG0158510 | Bifunctional inhibitor/lipid-transfer protein/seed storage 2S<br>albumin superfamily protein | -8.79 | 4.5E-05 |
| HORVU.MOREX.r2.4HG0288340 | translation initiation factor | -7.73 | 4.0E-07 |
| HORVU.MOREX.r2.7HG0561840 | MYB-related transcription factor | -7.69 | 4.0E-09 |
| HORVU.MOREX.r2.5HG0369380 | Glutaredoxin family protein | -7.29 | 1.0E-04 |
| HORVU.MOREX.r2.3HG0268450 | basic helix-loop-helix (bHLH)<br>DNA-binding superfamily protein | -6.98 | 2.3E-03 |
| HORVU.MOREX.r2.7HG0548630 | Pollen Ole e 1 allergen/extensin | -6.56 | 7.0E-09 |
| HORVU.MOREX.r2.5HG0356760 | Glutaredoxin family protein | -6.47 | 2.6E-03 |
| HORVU.MOREX.r2.5HG0412710 | CRT-binding factor | -6.45 | 1.5E-04 |
| HORVU.MOREX.r2.2HG0165830 | Pollen Ole e 1 allergen/extensin | -6.39 | 1.3E-09 |
| HORVU.MOREX.r2.4HG0330890 | Beta-xylosidase | -6.20 | 1.6E-03 |
| HORVU.MOREX.r2.3HG0268330 | dicer-like 3 | -5.83 | 4.2E-07 |
| HORVU.MOREX.r2.2HG0079590 | Avr9/Cf-9 rapidly elicited protein | -5.72 | 2.9E-03 |
| HORVU.MOREX.r2.7HG0623060 | Aquaporin | -5.61 | 4.4E-03 |
| HORVU.MOREX.r2.6HG0455680 | Pollen Ole e 1 allergen/extensin | -5.57 | 1.1E-07 |
| HORVU.MOREX.r2.5HG0438030 | Leucine-rich repeat receptor-like<br>protein kinase family protein | -5.47 | 1.3E-05 |
| HORVU.MOREX.r2.2HG0160110 | Orexin receptor type 2 | -5.39 | 1.0E-07 |

<sup>1</sup>FC= fold-change
<sup>2</sup>padj= adjusted *p* value

In the recovery samples, compared to the controls that never experienced drought, the most overexpressed gene was annotated as a leguminosin group485 secreted peptide, having 16-billion-fold more expression than in the controls (Table 3). This gene has been reported in *Medicago truncatula* root during root nodulation, with gene ID Medtr5g064530 and Medtr2g009450. However, a BLASTp of the transcript corresponding to the barley gene (HORVU.MOREX.r2.5HG0428280.1) has its best match (89% identity, e-value 2e-36) to a serine/threonine-protein phosphatase 7 (PP7) long-form homolog for barley (XP_044965344.1). Four genes showed more than 100-fold overexpression and 58 genes over 10-fold overexpression (Table S4). Compared to the drought samples, 139 genes showed more than 10-fold overexpression. The most overexpressed compared with drought (635 X; Table S7) was annotated as for a “bifunctional inhibitor/lipid-transfer protein/seed storage 2S albumin superfamily protein,” a protein type reported to have diverse functions (Wei & Zhong, 2014).

**Table 3.** Top 20 overexpressed genes during the recovery phase.

| Ensembl_gene_id | Description | log2 FC<sup>1</sup> | padj<sup>2</sup> |
| --- | --- | --- | --- |
| HORVU.MOREX.r2.5HG0428280 | Leguminosin group485 secreted peptide | 33.95 | 5.4E-10 |
| HORVU.MOREX.r2.5HG0406650 | Nucleolar-like protein | 7.59 | 2.4E-03 |
| HORVU.MOREX.r2.7HG0563500 | Fasciclin-like arabinogalactan protein | 7.11 | 2.7E-04 |
| HORVU.MOREX.r2HG0002090 | Arginine decarboxylase | 6.73 | 3.9E-03 |
| HORVU.MOREX.r2.7HG0604490 | Xyloglucan endotransglucosylase/hydrolase | 6.25 | 3.4E-03 |
| HORVU.MOREX.r2.2HG0083580 | Cytochrome P450 | 5.57 | 9.3E-03 |
| HORVU.MOREX.r2.4HG0340230 | translation initiation factor | 5.54 | 1.1E-03 |
| HORVU.MOREX.r2.2HG0179370 | Cytochrome P450 | 5.51 | 1.0E-02 |
| HORVU.MOREX.r2.7HG0579230 | MLG (mixed-linkage glucan) synthase, Biosynthesis of MLG (cell wall polysaccharide | 5.33 | 6.3E-04 |
| HORVU.MOREX.r2HG0072930 | Thiamine-phosphate synthase | 5.05 | 6.7E-15 |
| HORVU.MOREX.r2.6HG0519350 | Heparanase | 5.05 | 2.2E-04 |
| HORVU.MOREX.r2.2HG0141350 | 3-oxoacyl-[acyl-carrier-protein] synthase | 4.86 | 4.4E-08 |
| HORVU.MOREX.r2.3HG0183950 | Glycosyltransferase | 4.81 | 2.5E-03 |
| HORVU.MOREX.r2.5HG0355750 | Bidirectional sugar transporter SWEET | 4.80 | 3.5E-05 |
| HORVU.MOREX.r2.7HG0558300 | Glycosyltransferase | 4.78 | 4.4E-03 |
| HORVU.MOREX.r2.5HG0349900 | Receptor-like protein kinase | 4.67 | 2.4E-05 |
| HORVU.MOREX.r2.6HG0516480 | F-box protein SKIP8 | 4.52 | 1.5E-13 |
| HORVU.MOREX.r2.4HG0318550 | MYB transcription factor | 4.44 | 5.4E-03 |
| HORVU.MOREX.r2.6HG0502520 | 4-coumarate: coenzyme A ligase, Flavonoid biosynthesis | 4.39 | 7.9E-03 |
| HORVU.MOREX.r2.7HG0594280 | Beta-galactosidase | 4.33 | 6.7E-03 |

<sup>1</sup>FC= fold-change<sup>2</sup>padj= adjusted *p* value

The most underexpressed transcript in recovery compared to controls corresponded to a superfamily *Copia* retrotransposon, which automatic annotation designated as a “retrovirus-related Pol polyprotein”, having around 8-million-fold less expression than the control (Table 4). When drought and recovery DEGs were compared, the most strongly underexpressed (1.8-million fold) transcript likewise was annotated as a “Pol polyprotein from transposon *Tnt*1-94”, i.e., a superfamily *Copia* retrotransposon (Table S6). Of the top 20 repressed in recovery compared with drought samples (Table S6, three were annotated as dehydrins (435- to 1634-fold underexpressed) and three as heat-shock proteins (188- to 627-fold underexpressed). Compared to controls, overall, 3 genes showed more than 100-fold underexpression and 9 genes more than 10-fold underexpression (Table S5). Compared to drought samples, 222 genes in the recovery phase showed more than 10-fold underexpression and 140 showed more than 10-fold overexpression. To place the very many significantly differentially expressed genes into a biological context, we undertook a network analysis, as described below.

**Table 4.**
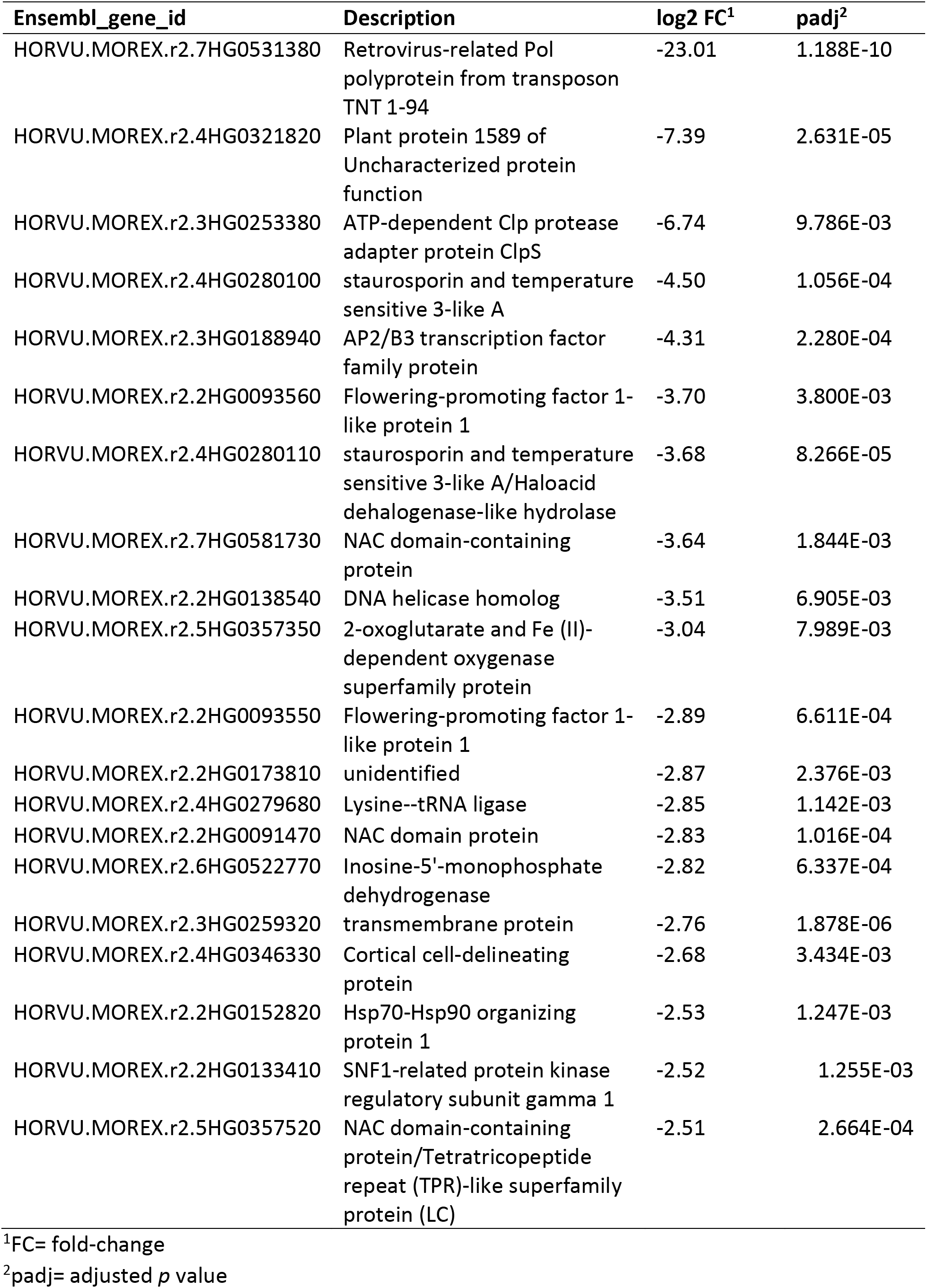
Top 20 underexpressed genes during the recovery phase.

### Network analysis

#### Upregulated networks under drought stress

When drought-treated plants were compared to the well-watered controls of the same age, a simple pattern of three clusters with two nodes, each comprising the upregulated genes, was seen (Fig. 7a). Detailed information on key nodes with the clusters is presented in Table S8. Cluster A, which has the highest statistical significance and contains 113 significantly upregulated genes, is annotated as “response to water” (GO:0009415) and to “acid chemical” (GO:0001101), meaning any process that responds to the availability of water (e.g., under drought), or to an anion. In this cluster, the dehydrin genes have the highest differential expression, ranging from 100- to 3458-fold overexpression during drought, followed by the late-embryogenesis-abundant (LEA) protein Lea14 at 60-fold (Table S12). Both are associated with physiological drought; dehydrin is a drought-inducible LEA II protein (Kosová *et al.*, 2014). Conversely, three dehydrins are among the top 20 underexpressed DEGs in recovery vs drought (Table S6). Of the other two clusters, one (cluster B) comprises nodes for response to salt stress (GO:0009651) and osmotic stress (GO:0006970), together with 126 significantly upregulated genes (Table S8). In cluster B, the most strongly upregulated genes are for a serine-threonine protein kinase (Table S12), which is a family well connected to drought response (Saddhe *et al.*, 2021) and salt stress (GO:0009651; 19-fold), and for an uncharacterized transcription factor (18-fold). Drought, salt, and osmotic stress response pathways are well-known to overlap (Buti *et al.*, 2019). The third cluster (C), with 250 genes upregulated during drought, is associated with negative regulation of protein metabolic processes in two nodes (GO:0051248, GO:0032269). These nodes contain multiple proteinase inhibitors, including a trypsin/amylase inhibitor such as was earlier shown to confer drought tolerance (Xiao *et al.*, 2013), that are strongly overexpressed in drought (64- to 694-fold).

**Fig. 7.**
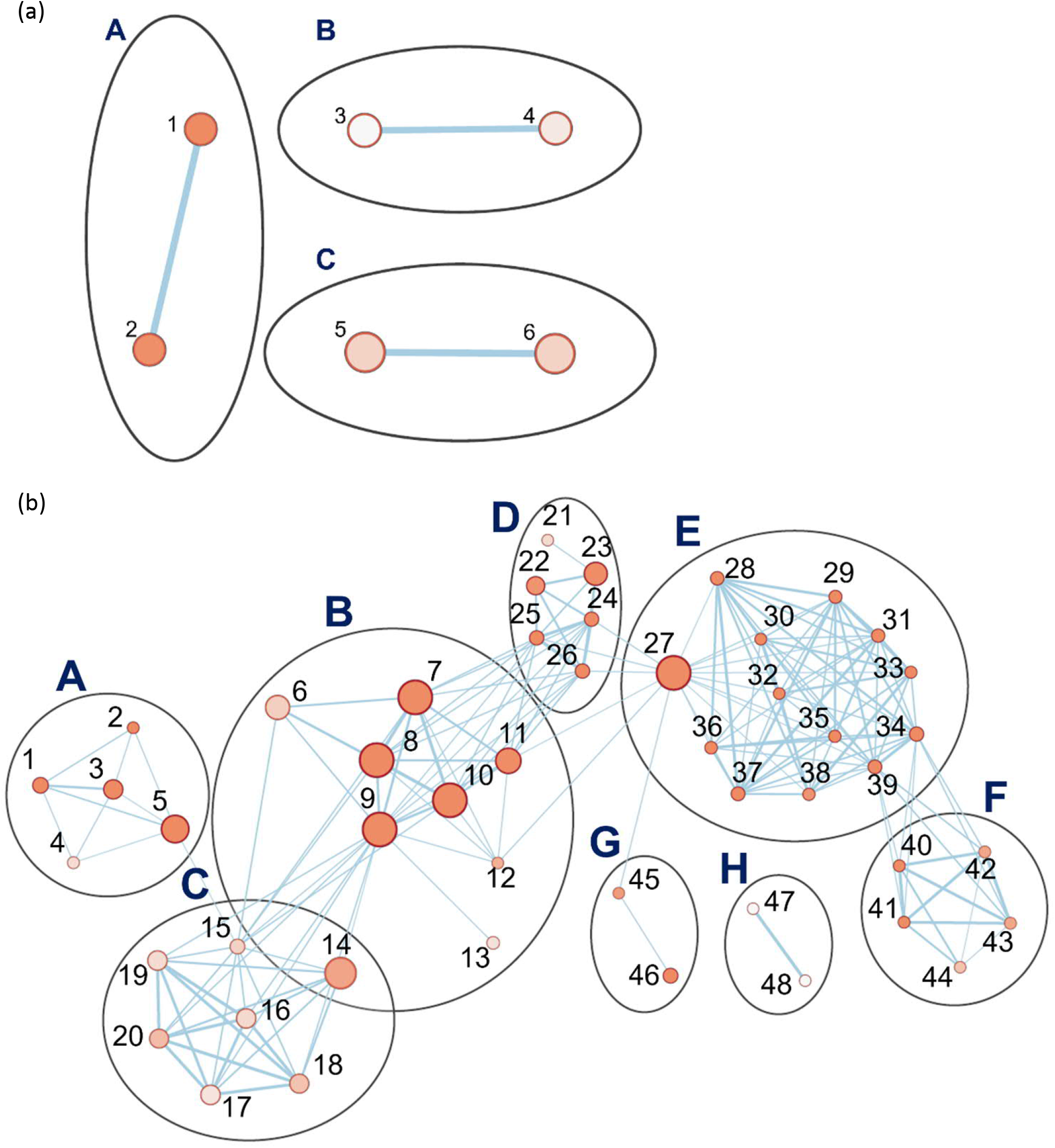
Pathway enrichment analysis of differentially expressed genes (DEGs) during the drought phase. (a) The set of three clusters of nodes containing upregulated genes (b) Eight interlinked clusters of interlinked nodes containing genes downregulated during the drought phase. Pathway nodes are shown as small circles connected by lines (edges) if the pathways share many genes. Nodes are colored by enrichment score, and the thickness of lines according to the number of genes shared by the connected pathways. Nodes are numbered; descriptions for upregulated and downregulated genes are organized by node number in Tables S6 and S7, respectively.

#### Downregulated networks under drought

During the drought stress, 8 clusters, comprising 48 nodes, were downregulated (Fig. 7b). All are related to processes of growth and development. The cluster with the highest significance and with the greatest number of genes (B) is related to metabolism of carboxylic acids and anions and comprises 3600 DEGs. These are involved in nutrient supply, plant development, and processes including cell wall signaling, guard mother cell division, and pathogen virulence (Polko & Kieber, 2019). The key nodes in this cluster include metabolism of organic acids (GO:0006082), oxoacids (GO:0043436), cellular amino acids (GO:0006520), carboxylic acids (GO:0019752), and of small molecules (GO:0044281) (Table S9). The other interconnected clusters are associated with cluster A (654 genes), which concerns the regulation of photosynthesis, (e.g., photosystem genes, 2- to 5-fold underexpressed): purine ribonucleotide metabolism, tRNA aminoacylation translation, pigment biosynthesis and tetraterpenoid biosynthesis. A second set of linked clusters downregulated under drought stress comprises plastid translation organization (Cluster G, 83 genes; GO:00032544, GO:0009657) and pigment, including chlorophyll and carotenoid, biosynthesis (Cluster E, 1588 genes; Cluster F, 136 genes). A small cluster (H, 10 genes) unlinked to others, is annotated as concerning plant ovary development (GO:0032544) though, given leaf samples were analyzed, the function appears to be otherwise. Detailed information on the key nodes and clusters can be found in Tables S9 and S13. Hence, the downregulated pathways are manifold and generally involved in growth and development.

#### Upregulated networks during recovery

Recovery is the process of restoring the physiological and molecular functions from drought-induced damage (Chen *et al.*, 2021). As described above, gene networks related to the growth and development were downregulated during drought. In turn, during recovery, many associated with growth, plastidial function, and energy metabolism were upregulated, organized in five linked clusters of interconnected nodes (Fig. 8a). Cluster A had 2597 genes significantly upregulated and concerns cell wall metabolism (Table S10). Its nodes include those for carbohydrate (GO:0005975), polysaccharide (GO:0005976), glucan (GO:0006073), and xyloglucan (GO:0010411) metabolism, “cell wall organization or biogenesis”, “external encapsulation structure organization” (GO:0045229), and hemicellulose metabolism (GO:0010410). Genes for biosynthesis of xyloglucan, an abundant component of the primary cell wall, were particularly strongly induced, up to 239-fold over the control plants that did not experience drought. Two of the clusters (B, 2522 genes upregulated; C, 7842 genes) include nodes associated with protein biosynthesis that were downregulated during drought but are upregulated during recovery. These include nodes for ribosome biogenesis (GO:0042254), ncRNA metabolism (GO:0034660), rRNA processing (GO:0006364), ribonucleoprotein complex assembly (GO:0022618), maturation of LSU-rRNA (GO:0000463), translation (GO:0006412), peptide biosynthesis and metabolism (GO:0043043, GO:0006518), amide biosynthesis (GO:0043604), and tRNA aminoacylation and metabolism (GO:0043039, GO:0006399). Consistent with reactivation of leaf function, plastid, and chloroplast organization processes (GO:0009657, GO:0009658) were upregulated. One unlinked cluster, for “quinone process” (GO:1901663), nevertheless also indicates upregulation of photosynthetic function and growth, as plastoquinone and ubiquinone serve in the electron transport chains respectively of photosynthesis and aerobic respiration. Descriptions of the clusters and their nodes are found in Tables S10 and S14.

**Fig. 8.**
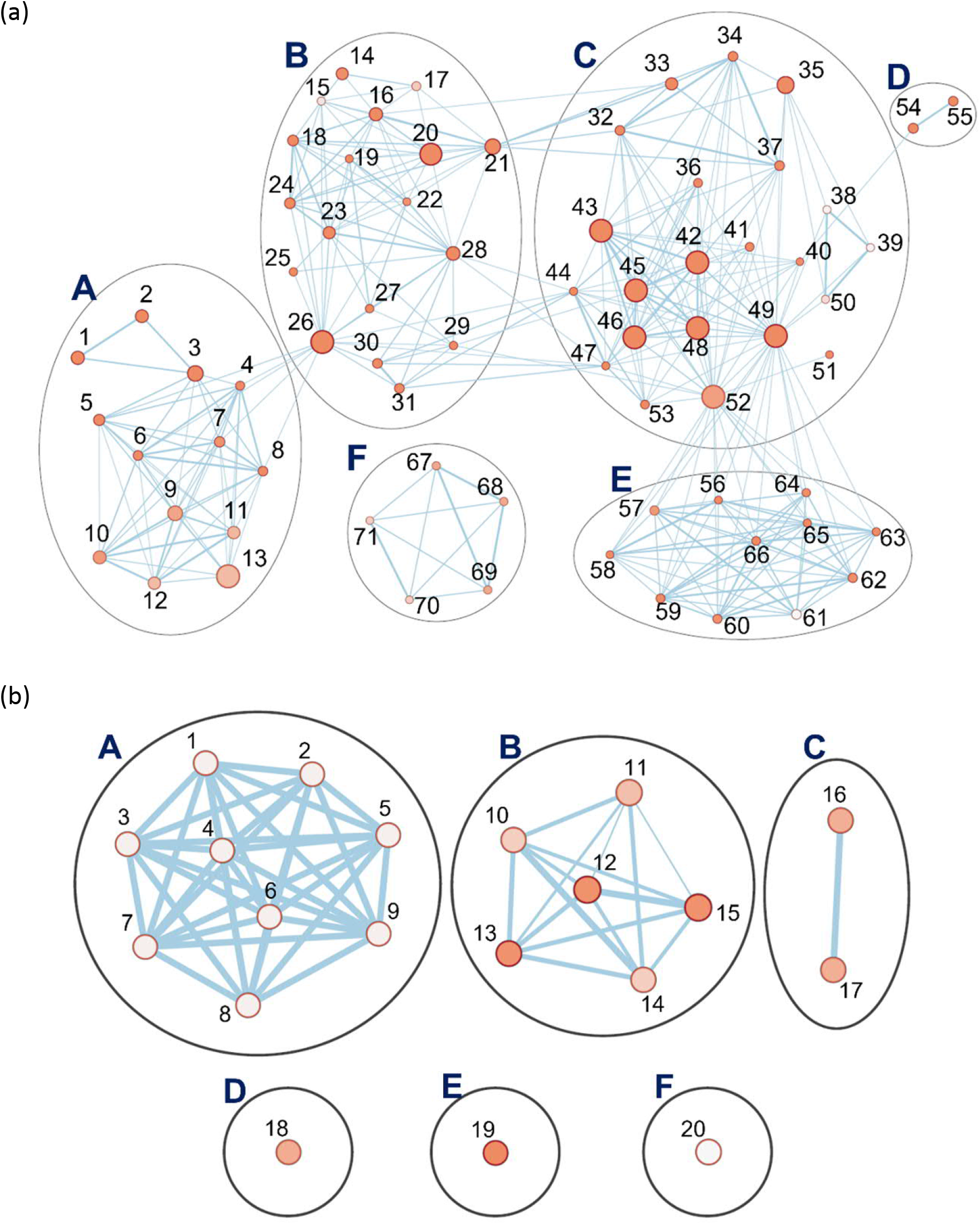
Pathway enrichment analysis of differentially expressed genes (DEGs) during the recovery phase. (a) Six interlinked clusters of interlinked nodes containing upregulated genes (b) Six clusters of nodes containing downregulated genes. Pathway nodes are shown as small circles connected by lines (edges) if the pathways share many genes. Nodes are colored by enrichment score, and the thickness of lines according to the number of genes shared by the connected pathways. Nodes are numbered; descriptions for upregulated and downregulated genes are organized by node number in Table S8 and S9, respectively.

#### Downregulated networks during recovery

The networks of genes downregulated during recovery, when compared to the controls, form six clusters (Fig. 8b). Cluster F (13 genes in one node) is associated with the de-repression of genes associated with organelle organization (GO:0010638). Cluster A has nine nodes, each with a single gene, for processes spanning cell wall macromolecules (GO:0010981), carbohydrate metabolism (GO:0010676), trichome papilla formation (GO:1905499), among others. Descriptions of the clusters and their nodes are presented in Tables S11 and S15.

Of the other five clusters of genes downregulated during recovery, notably, three (B, D, E) contain altogether 117 networked autophagy-related (ATG) genes involved in ubiquitin-mediated autophagy (Fig. 8b). The largest cluster (B), has six nodes, including ones with genes concerned with autophagosome organization and assembly. Two unlinked ATG clusters (D, E) consist of a single node each. One is for genes involved in protein K63-linked de-ubiquitination (GO:0070536). In this cluster, the genes for deubiquitinating enzyme AMSH1 are also observed. The other cluster with its single node concerns genes involved in protein neddylation (GO:0045116), the process by which the ubiquitin-like protein NEDD8 is conjugated to its target proteins in a manner analogous to ubiquitination.

The reduction in expression of the ATG genes is modest, compared with the controls, around two-fold at most. However, of the six gene clusters with reduced expression during recovery, the top three most strongly reduced are all for ATG genes. The *p-*values of the ATG nodes are all highly significant (mean 0.01). Among the ATG genes most strongly downregulated: autophagy-related protein 101 (−1.76X), beclin 1 (−1.7 X), autophagy-related protein 3 (−1.48X), all in cluster B (autophagy); a 26S proteasome regulatory subunit (−1.98X) in cluster D (protein K63-linked deubiquitination); ubiquitin-conjugating enzyme E2 (−1.57 X), in cluster E (neddylation). When gene expression levels at recovery is compared to that at drought, a similar picture emerges for ATG genes (Table S15). While only a few are significant at padj = 0.05, most show the same trends.

### *BARE* response during drought and recovery

For all four varieties during drought and recovery (Tables 5, S16), the mRNA levels matching the *BARE gag* region was analyzed by qPCR; in GP, additionally the abundance of mRNA containing the 5’ UTL regions corresponding to expression from TATA1 and TATA2 (Chang & Schulman, 2008), respectively for the gRNA and mRNA, was measured (Table 5). As an indicator of general stress response, *hsp17* expression was examined; putative *hsp17* and *hsp21* were among the most strongly downregulated genes in recovery vs drought (Table S6).

**Table 5.** Relative expression of *hsp17* and *BARE* transcripts in Golden Promise during the drought and recovery phases along with TATA1 and TATA2 –driven mRNA levels.

| DROUGHT |  |  |  |  |
| --- | --- | --- | --- | --- |
| Samples | *hsp17* ** | *gag* ** | TATA1 ** | TATA2 ** |
| Control 1 | 1.50 ± 0.08 | 0.66 ± 0.17 | 0.49 ± 0.09 | 0.60 ± 0.11 |
| Control 2 | 2.75 ± 0.38 | 1.19 ± 0.09 | 0.83 ± 0.05 | 0.40 ± 0.01 |
| Drought 1 | 8.01 ± 1.96 | 4.78 ± 0.30 | 5.15 ± 0.82 | 1.78 ± 0.24 |
| Drought 2 | 4.06 ± 0.10 | 3.52 ± 0.79 | 2.12 ± 0.20 | 0.87 ± 0.19 |
| RECOVERY |  |  |  |  |
| Samples | *hsp17* | *gag* | TATA1 * | TATA2 |
| Control 1 | 0.64 ± 0.03 | 1.17 ± 0.07 | 0.57 ± 0.08 | 1.03 ± 0.13 |
| Control 2 | 0.39 ± 0.09 | 0.35 ± 0.14 | 1.33 ± 0.12 | 0.59 ± 0.04 |
| Recovery 1 | 0.38 ± 0.05 | 0.36 ± 0.11 | 0.23 ± 0.01 | 0.97 ± 0.08 |
| Recovery 2 | 0.37 ± 0.02 | 0.73 ± 0.11 | 0.61 ± 0.07 | 1.06 ± 0.21 |

Values are given as means ± SD of three technical replicates for each plant. The asterisks indicate the significance level of \* $p \leq 0.05$ , \*\* $p < 0.005$ analysed using a non-parametric Mann Whitney U-test.

Drought induced expression of both *hsp17* and of *BARE* (*gag*) was significantly higher (Tables S17, S18) in GP and Arvo over the well-watered control (Table 5, S16). During recovery, *BARE gag* and *hsp17* were downregulated in Arvo and GP (down 2.2 X, Arvo; 1.3 X, GP), but not in H673 (up 2.9 X over control), suggesting delayed recovery in H673 (Table 5, S16). Strikingly, in Morex, *gag* levels but not *hsp17* showed an inverse pattern from the other lines: decreasing 1.8-fold under drought but increasing 1.7-fold during recovery (Table S16). We additionally followed the expression of the 5’ UTLs corresponding to both the gRNA (TATA1) and the mRNA (TATA2) of *BARE* in GP. TATA1 was more strongly upregulated than TATA2 under drought (Tables 5, S17). During recovery, TATA1 and HSP17 expression were below that of the well-watered control, whereas TATA2 was near to control levels. On the protein level (Fig. S5), *BARE* Pol, produced from the TATA2 mRNA, showed a strong drought response, which decreased but was still above the control during recovery.

## DISCUSSION

Drought tolerance and resilience are important traits for wild plants and commonly sought for crops. For crops, in contrast to wild species, not only the production of viable seed, but also high yield, is desirable. Hence, trade-offs between drought tolerance and concomitant stomatal water loss and CO_2_ uptake are relevant to breeding. Drought under field conditions is complex, affected by many factors: ambient heat, humidity, VPD, and their diurnal fluctuations; soil structure and hydration profile; soil and root microbiomes; the length and completeness of rain cessation (Scharwies & Dinneny, 2019). Likewise, experimental droughts in greenhouses and growth chambers can differ greatly from each other and from those in the field. Here, we sought to correlate transcriptional to physiological responses in barley on a precision feedback lysimeter platform with well-controlled soil and watering as well as continuous data collection. We recorded VPD and PAR continuously throughout the experiment, as well as the daily transpiration and increase in plant weight. The platform itself was in a greenhouse that closely reflected natural external environmental conditions beyond.

### Physiological response of four barley varieties to drought and rewatering

We analyzed the drought and recovery responses of four barley varieties for which pilot experiments showed a difference in transpiration rates and ϴ_crit_. We have not tracked leaf water potential, but rather the stomatal response to SWC; we use the terms anisohydric and isohydric as equivalent to ϴ_crit_ at comparatively low and high SWC, respectively. The SWC at which drought response is invoked is genotype-dependent; the two strategies can be found within genotypes of the same species such as barley or wheat (*Triticum aestivum* L.), which have both spring- and autumn-sown types that thereby experience contrasting rainfall patterns (Tao *et al.*, 2017) depending on the cultivation region (Galle *et al.*, 2013).

Morex, having the highest daily transpiration rate, reached ϴ_crit_ at VWC 0.46 cm^3^/cm^3^ on day 36, whereas GP reached ϴ_crit_ at the lowest VWC, 0.42 cm^3^/cm^3^, already on day 33 and continued to dehydrate at the fastest rate, but with the lowest initial transpiration rate. Hence, the rate of water loss after ϴ_crit_ was not links to the initial transpiration rate, as GP dried the fastest and Arvo the slowest. Our interpretation is that GP is therefore the most anisohydric of the four. Bred for the UK, GP may be able to manage the risk as SWC falls, due to its low transpiration rate and the relatively balanced precipitation throughout the growing season. Morex also reaches ϴ_crit_ at a comparatively low VWC but, nevertheless, maintains weight as well as H673, which has ϴ_crit_ at the highest VWC, so would appear to manage water loss well. Arvo appears to be the most isohydric; it, like H673, responds at a high ϴ_crit_ and maintains its weight best during drought. This is consistent with Arvo and H673 being varieties from Finland, where droughts occur early in the growing season and can persist for weeks, but where the probability of precipitation increases as the season progresses (Tao *et al.*, 2017). All lines responded to rewatering by increasing specific weight gain (g/g), GP being the lowest, but none returned to control rates during the three days that recovery was followed. Full recovery may require longer.

### Gene networks and differential expression in Golden Promise during drought and recovery

We focused gene expression analyses on GP, as the standard line for transformation and editing. *Drought*: Networks of genes responding to drought, salt, and acid, as well as negative regulators of metabolism, were upregulated following ϴ_crit_. Among individual genes, *glutathione s-transferase* (*GST*) was most upregulated (3 x 10^6^ -fold) over the control; at the sampling point early in recovery, it had dropped by 8.6-fold relative to drought samples. GST expression is associated with drought tolerance in barley (Rezaei *et al.*, 2013); it scavenges drought-produced ROS for detoxification (Guo *et al.*, 2009). Likewise, increased GST expression was observed under cold and osmotic stress in a potato genotype tolerant to these (Guo *et al.*, 2009). The second most highly upregulated (447 X) gene is chloroplastic linalool synthase (XP_044955412), a terpenoid biosynthetic enzyme; its expression fell 70-fold in recovery relative to drought. In rice, the bHLH family transcription factor *RERJ1* is drought-induced, leading to jasmonate (JA) biosynthesis and induction of linalool synthase (Valea *et al.*, 2022). In barley, 98 among 135 detected phenolic and terpenoid compounds (linolool was not investigated) were shown to change in expression as result of drought stress (Piasecka *et al.*, 2017). Our observed induction of linolool synthase here is consistent with a role in ABA/ JA-mediated stomatal closure as reported for Arabidopsis (Munemasa *et al.*, 2019). *Dehydrin* (*Dhn*) transcripts were also highly upregulated, 3500 X over control, and among the most strongly downregulated in recovery vs drought. Dehydrins, along with other late embryogenesis-abundant (LEA) proteins such as LEA14 (also upregulated here), protect enzymes from dehydration and other environmental stresses (Liu *et al.*, 2017). Differential *Dhn* expression was shown in cultivated and wild (*H. spontaneum*) barley to be correlated with drought tolerance (Suprunova *et al.*, 2004; Kosová *et al.*, 2014), likewise in *Brachypodium distachyon* (Decena *et al.*, 2021), wheat (Brini *et al.*, 2007; Wang *et al.*, 2014) and rice (Verma *et al.*, 2017).

Networks for photosynthetic and plastidial processes, metabolism, and ovary development were downregulated during drought, consistent with earlier reports for some individual genes from these processes: ribulose 1,5-bisphosphate carboxylase small subunit (*rbcS*), chlorophyll a/b-binding protein (*cab*), and components of photosystems I and II (Seki *et al.*, 2002; Narusaka *et al.*, 2004). Here, the most strongly underexpressed gene was for cytochrome P450 (*CYP*), which conversely was the third most strongly overexpressed in recovery vs drought. The *CYP* genes are widely distributed in plants and animals (Shiota *et al.*, 2000), forming a large family, 45 in wheat (Li & Wei, 2020), playing multiple roles (McKinnon *et al.*, 2008), including in biotic and abiotic stress response (Li & Wei, 2020). In rice, 83 *OsCYPs* are differentially expressed, of which 6 are strongly induced and 3 strongly downregulated, during drought (Wei & Chen, 2018).

Two genes for cortical cell delineating protein were highly (−705 X, - 621 X) repressed during drought; while associated with roots, they appear to have a role in cell division and elongation (Isayenkov *et al.*, 2020). These, too, had their expression strongly upregulated during recovery vs drought. Downregulated almost as strongly (−442 X) was a gene of the *bHLH* family. This widespread family of transcription factors has been shown in various plants, including wheat, to be integrated into ABA- and JA-mediated signaling, ROS scavenging, and stomatal closure, as well as to play multiple roles in development (Yang *et al.*, 2016; Guo *et al.*, 2021). Given that *bHLH* is strongly induced by drought and ABA, the pattern seen here suggests that induction occurs before ϴ_crit_ and that its downregulation thereafter may be linked to suppression of developmental processes.

#### Recovery

During the recovery phase, gene networks connected to growth and to chloroplast development were upregulated; these processes were downregulated under drought. The putative *PP7*, here the most upregulated individual gene vs control during recovery, has shown to be highly expressed in Arabidopsis stomata (Andreeva *et al.*, 1999), to regulate phytochrome signaling (Andreeva *et al.*, 1999) and to be involved also in Arabidopsis chloroplast development (Xu *et al.*, 2019), a function that would be consonant with recovery in the barley system.

Notably, autophagy processes were downregulated. Autophagy mechanisms are conserved in eukaryotes for turning over unwanted cytoplasmic components and maintaining homeostasis during stress (Marshall & Vierstra, 2018; Bao, 2020; Tang & Bassham, 2022). Mutant or silenced individual autophagy genes (*atg5, ATG6, atg7*, *ATG8d, ATG18h*), variously in Arabidopsis, tomato, and wheat (Liu *et al.*, 2009; Marshall & Vierstra, 2018; Zhu *et al.*, 2018; Li *et al.*, 2019), reduced drought tolerance. In barley, a mutant of autophagosome formation gene *ATG6* (*Beclin 1*) was upregulated under various abiotic stresses including drought, whereas its knockdown resulted in yellowing leaves in dark and H_2_O_2_ treatments (Zeng *et al.*, 2017). We confirmed that *ATG6* is identical to 3HG0280440.1 (Morex assembly vers3) and saw a 1.47 X reduction in recovery vs drought (adjusted *p*=0.01). While these studies indicated the importance of individual autophagy components in mediating abiotic stress response including drought, the autophagy network as a whole had not been examined. During recovery, along with *ATG6*, we observed a downregulation of many other *ATG*s in nodes connected either to initiation of autophagy, vesicle nucleation, expansion, or autophagosome formation. The likely importance of the suppression of autophagy during recovery is indicated by COST1 (constitutively stressed 1), attenuating autophagy in Arabidopsis under optimal growth conditions; *cost1* mutants are highly drought tolerant but very small, growing slowly (Bao *et al.*, 2020). Of the two *COST1* matches in the barley genome, expression of HORVU.MOREX.r2.4HG0296600.1 was not seen; for HORVU.MOREX.r2.6HG0480550, expression was almost identical in drought and recovery samples and regarding the controls.

### Response of retrotransposon *BARE* to drought and recovery

*BARE*, of which the reference genome contains 20,258 full-length copies and 6216 solo LTRs (Mascher *et al.*, 2021), carries out the various stages of the retrotransposon lifecycle: it is transcribed, translated, and forms virus-like particles (VLPs) (Jääskeläinen *et al.*, 2013). While the *BARE* LTR promoter was earlier shown to carry ABA response elements (ABREs) (Suoniemi *et al.*, 1996), the capsid protein Gag to be more strongly expressed under drought (Jääskeläinen *et al.*, 2013), and *BARE* copy number variations and insertional polymorphism patterns to be consistent with long-term drought activation (Kalendar *et al.*, 2000), *BARE* transcription has not earlier been linked to physiologically characterized drought. Here, we found that *BARE* transcription, as measured by qPCR from the *gag* coding region, increased following ϴ_crit_ in GP, Arvo, and H673, as did the gene for heat-shock protein *HSP17,* a marker for drought stress (Guo *et al.*, 2009); Morex, however, showed *HSP17* response but no increased *BARE* transcription. For Arvo and GP, both the *BARE gag* and *HSP17* were strongly lower in the samples taken after rewatering, whereas H673 had apparently not entered recovery, given that the *HSP17* level was still elevated. Divergent *BARE* levels in different varieties are consistent with those seen earlier (Jääskeläinen *et al.*, 2013), where GP showed a stronger response on the Gag protein level than did cv. Bomi. For GP, we also examined the drought response for the two transcripts, one driven by TATA1, which produces the gRNA that will be reverse transcribed, and another by TATA2, which produces the translatable mRNA. In drought, TATA1 was more strongly induced than TATA2, whereas during recovery, TATA1 showed lower expression than in control samples, perhaps indicating post-transcriptional silencing of the *BARE* gRNA. Consonant with the qPCR results, the most strongly downregulated gene in the recovery RNA-seq (HORVU.MOREX.r2.7HG0531380) matches best a superfamily *Copia* integrase domain (KAE8773625.1; 65% similar residues, e 7 x 10^-101^), likely from the *Hopscotch* family.

### Conclusions

Using a precision phenotyping platform, we have identified barley varieties whose transpiration rates drop (ϴ_crit_) at different points as soil dries during drought, which may mirror adaptation to pre-flowering vs terminal drought, or to droughts of varying length. The timing of droughts regarding phenology are expected to shift across Europe over coming decades (Appiah *et al.*, 2023). Using ϴ_crit_ as a guidepost, we carried out a global analysis of gene expression in the transformable variety GP in response to drought and rewatering and placed the DEGs into the context of biological pathways. We identified several strongly differentially expressed genes not earlier associated with drought response in barley, including for linalool synthase, cortical cell delineating protein, and long-form PP7. These data will serve to establish candidate genes from phenotyped mapping populations and diversity sets and for analysis of the many drought tolerance QTLs heretofore identified (Zhang *et al.*, 2017; Moualeu-Ngangue *et al.*, 2020). We were able to confirm that the *BARE* retrotransposon family is strongly transcriptionally upregulated by drought and downregulated during recovery unequally between cultivars and that the two *BARE* promoters are differentially drought responsive. In the context of recovery rate and resilience, which is key to maintenance of crop yields in the face of climate change, our observation that the autophagy gene network is downregulated during recovery is worthy of further investigation; we are currently constructing a higher resolution timeline.

## Data availability

The data that support the findings of this study can be found can be found in the EBI Array Express database as accession E-MTAB-12732, or within the supplementary material of this article, or are available from the corresponding author upon reasonable request.

## Acknowledgements

We acknowledge Anne-Mari Narvanto for excellent technical assistance and Triin Vahisalu for useful discussions on the physiology and molecular biology of drought response. Funding was provided by the Finnish Ministry of Agriculture and Forestry within the ERA-NET SusCrop ClimBar and BARISTA projects and by the Academy of Finland project “Retrostress” under Decision 314961.

## Author contributions

M.P. carried out the physiological experiments and expression assays, analyzed the data, and wrote the manuscript; J.T. was responsible for bioinformatics analyses; MJ carried out protein expression analyses; WC contributed to qPCR analyses; A.D. contributed to physiological experiments; M.M. and A.H.S. designed the experiments; A.H.S. revised the manuscript and was project P.I.

## Supporting Information

### Supplementary Figures

**Fig. S1.**
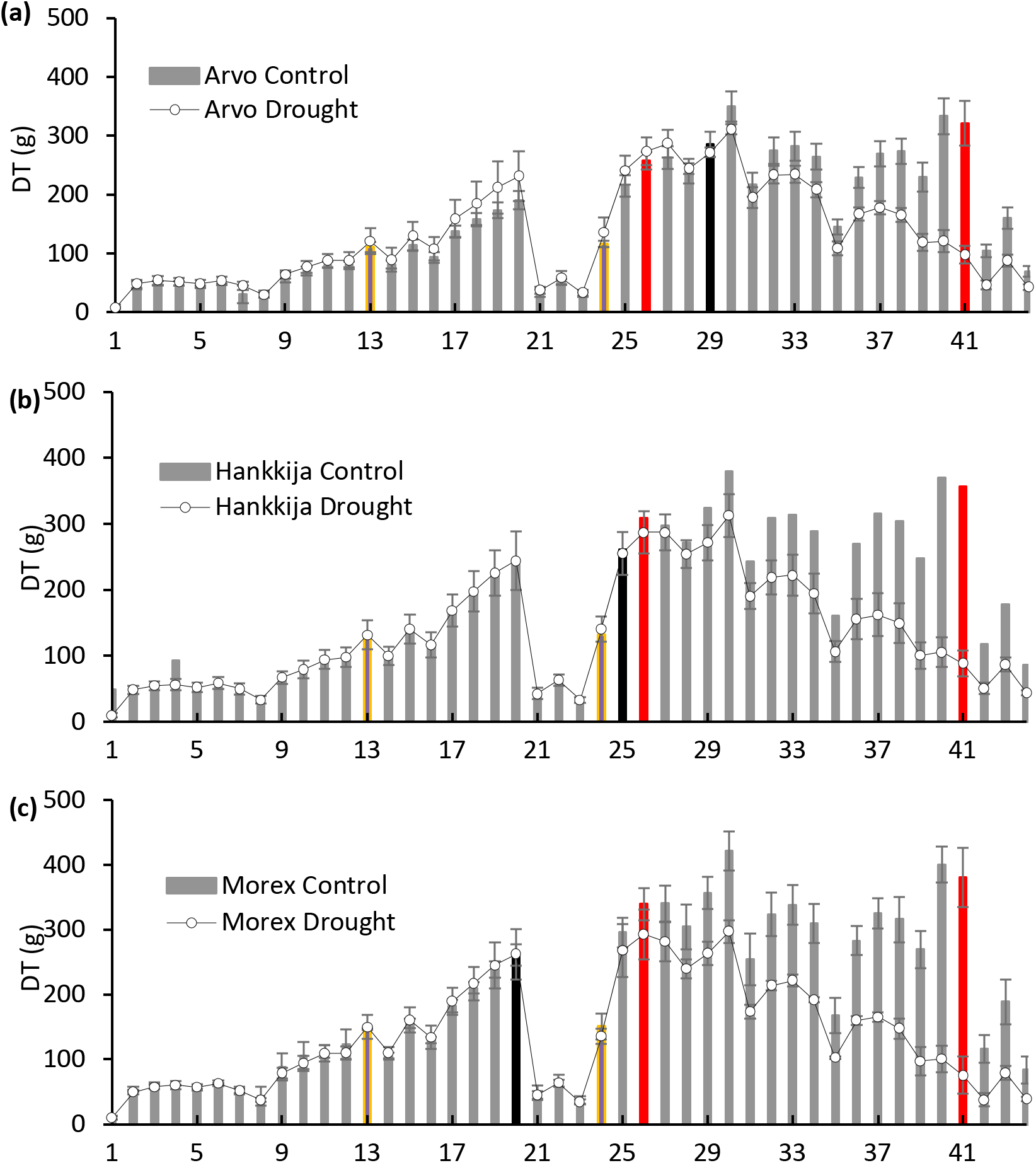
Daily transpiration. (a) Arvo, (b) Hankkija 973 (H973), (c) Morex. Yellow bars represent the start and end of Dry Phase I, red bars the start and end of Dry Phase II. The black bar represents the day when the transpiration of the droughted plants diverged from the controls.

**Fig. S2.**
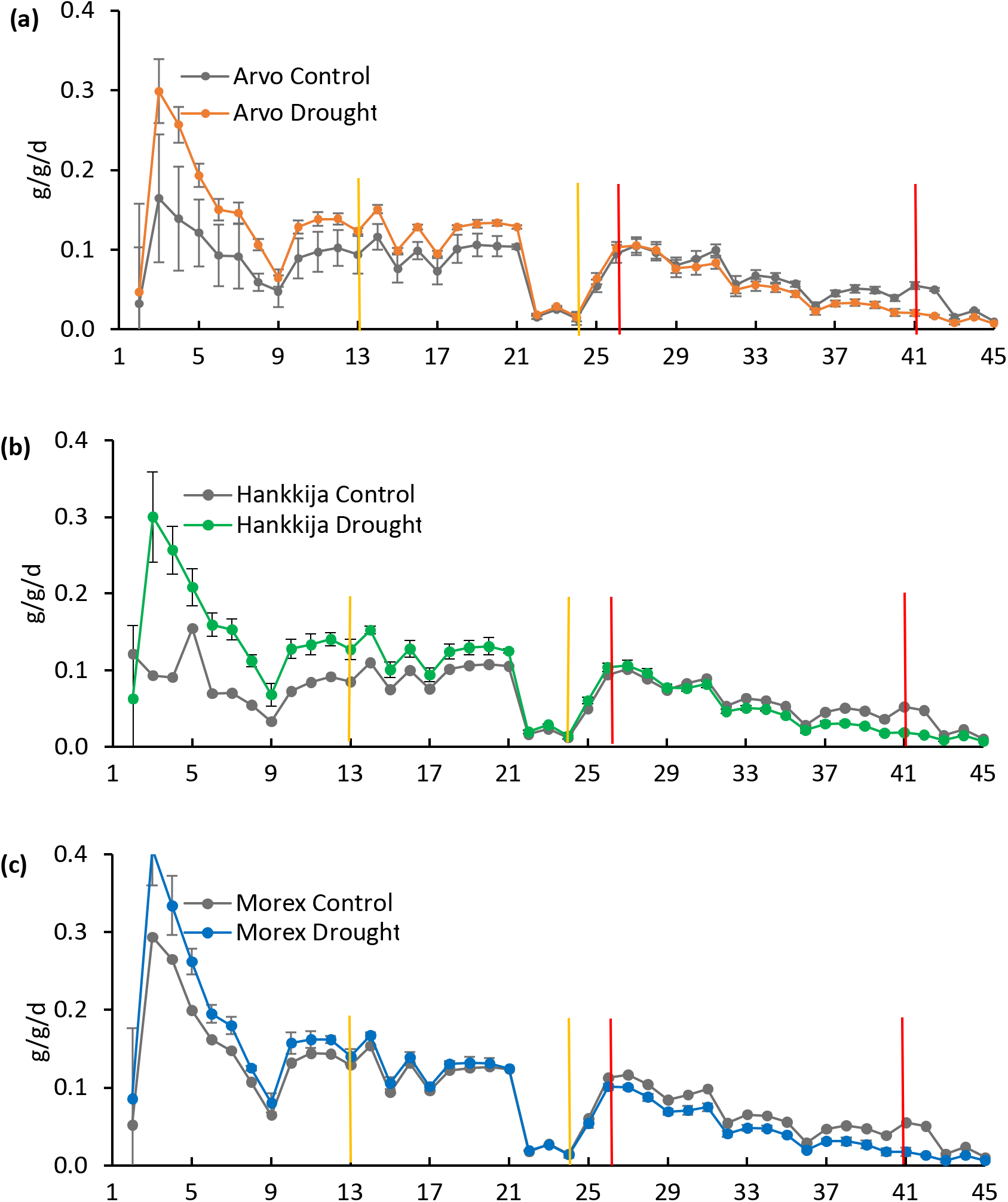
Daily change in specific plant weight. (a) Arvo, (b) Hankkija 973 (H973), (c) Morex. Yellow bars represent the start and end of Dry Phase I, red bars the start and end of Dry Phase II.

**Fig S3.**
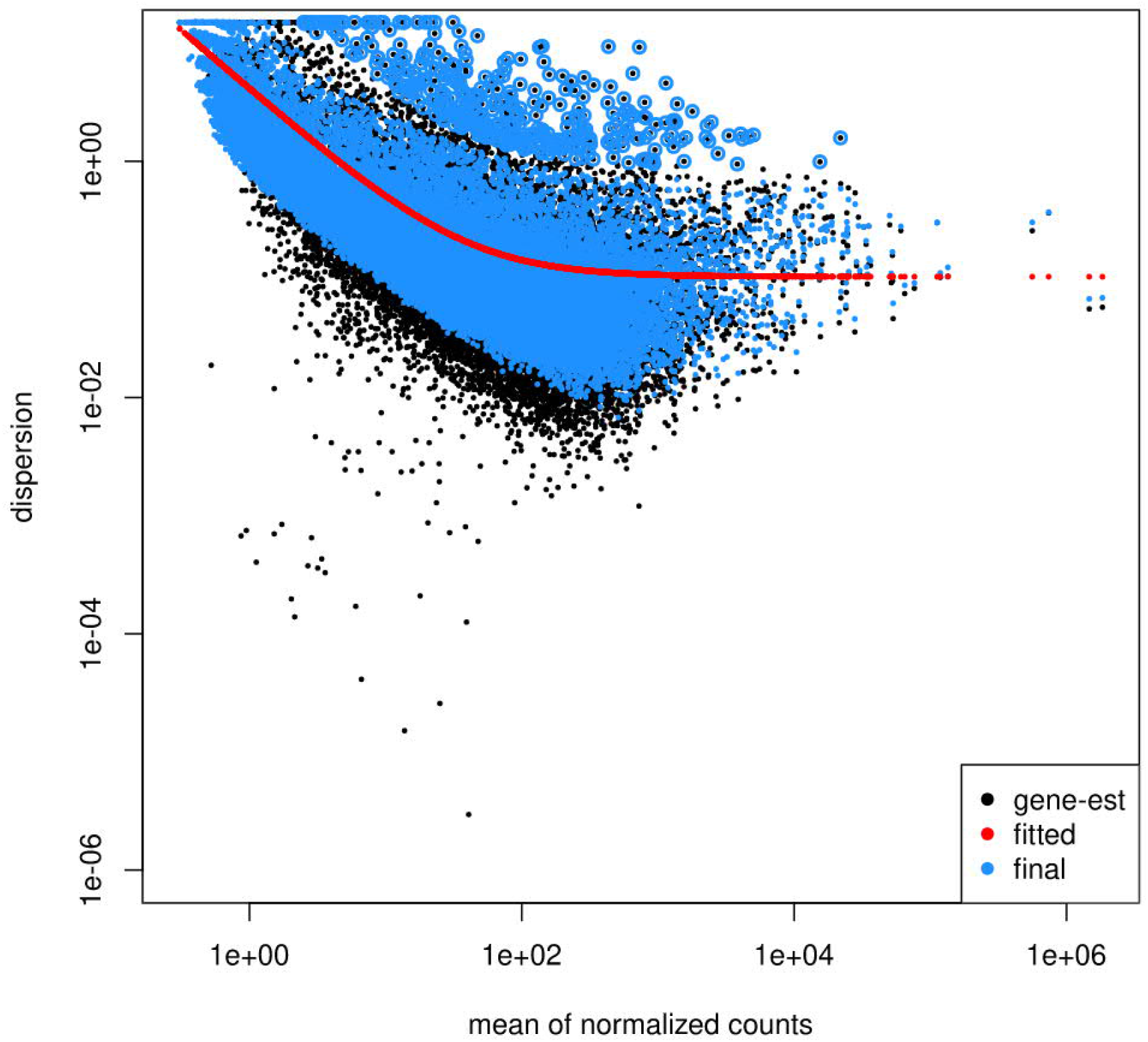
Plot of dispersion estimates. DESeq2 dispersion plot showing dispersion of each gene (black), the trend line for all samples (red), the corrected value of dispersion (blue), and outliers (black dot surrounded in blue) are shown.

**Fig. S4.**
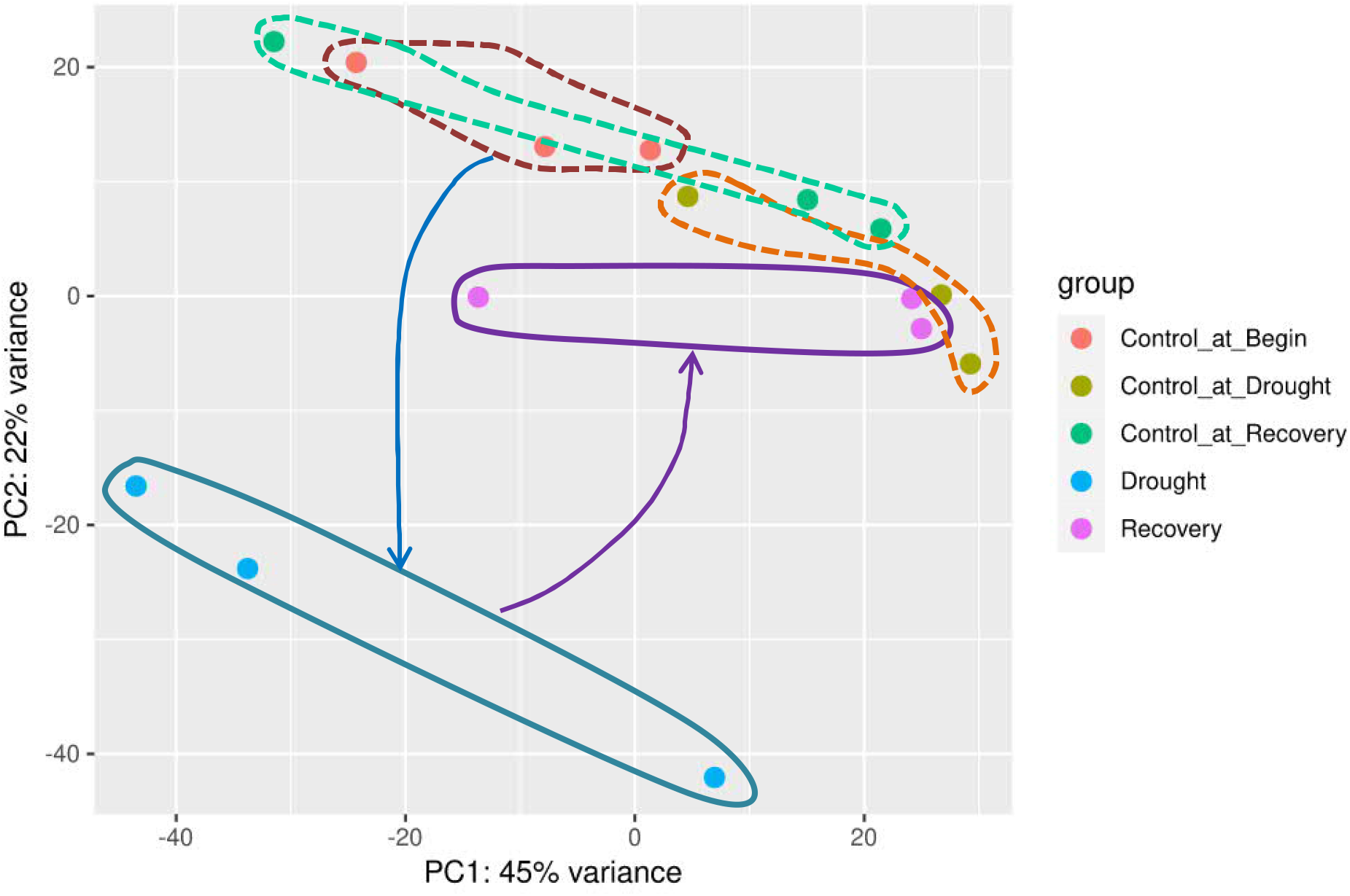
Principal Components Analysis (PCA) plot generated from RNA-seq data. DeSeq2 plot of the two major variance components for the groups of samples. Groups are differentiated by different colors: control at beginning (orange), control at drought (olive green), control at recovery (green), drought (blue), and recovery (purple). Samples within each group are bounded by lines in the group color, dashed for the controls. The arrows suggest the movement of gene expression pattern through two-dimensional variance space during the course of the experiment.

**Fig. S5.**
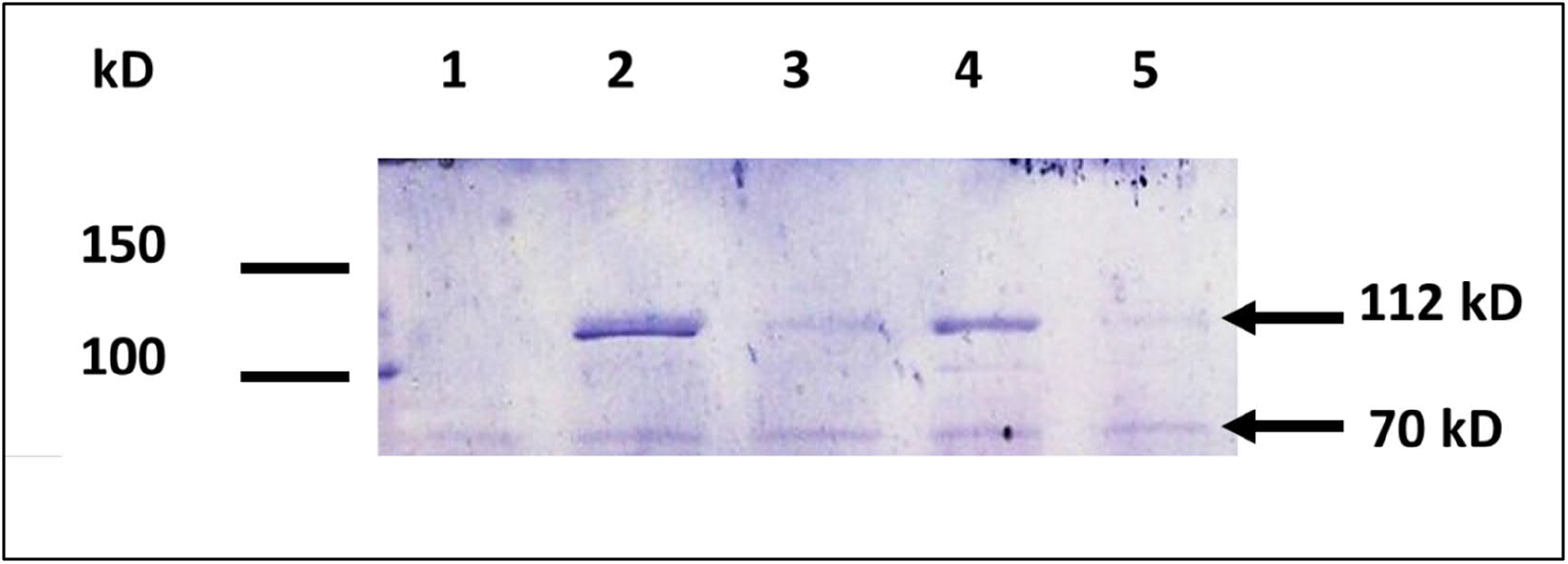
Comparison of BARE Pol protein levels in drought and recovery over control in GP. (1) day 12, well-watered control; (2) day 41, drought; (3) day 41, control; (4) day 44, rewatered; (5) day 44, control.

### Supplementary Tables (in excel file)

Table S1. Primers used in qPCR.

Table S2. Overexpressed genes, drought phase.

Table S3. Underexpressed genes, drought phase.

Table S4. Overexpressed genes, recovery phase.

Table S5. Underexpressed genes, recovery phase.

Table S6. Top 20 underexpressed genes, recovery vs drought.

Table S7. Top 20 overexpressed genes, recovery vs drought.

Table S8. Network analysis of overexpressed genes, drought phase.

Table S9. Network analysis of underexpressed genes, drought phase.

Table S10. Network analysis of overexpressed genes, recovery phase.

Table S11. Network analysis of underexpressed genes, recovery phase.

Table S12. Clusters and representative network nodes, overexpressed genes, drought phase.

Table S13. Clusters and representative network nodes, underexpressed genes, drought phase.

Table S14. Clusters and representative network nodes, overexpressed genes, recovery phase.

Table S15. Clusters and representative network nodes, underexpressed genes, recovery phase.

Table S16. Relative expression of *HSP17* and *BARE gag* mRNA levels.

Table S17. Mean rank by Mann Whitney U test of qPCR expression levels for GP.

Table S18. Mean rank by Mann Whitney U test of pPCR expression levels for Arvo, H673, and Morex.

